# The *Solanum tuberosum GBSSI* gene: a target for assessing gene and base editing in tetraploid potato

**DOI:** 10.1101/628107

**Authors:** Florian Veillet, Laura Chauvin, Marie-Paule Kermarrec, François Sevestre, Mathilde Merrer, Zoé Terret, Nicolas Szydlowski, Pierre Devaux, Jean-Luc Gallois, Jean-Eric Chauvin

## Abstract

Genome editing has recently become a method of choice for basic research and functional genomics, and holds great potential for molecular plant breeding applications. The powerful CRISPR-Cas9 system that typically produces double-strand DNA breaks is mainly used to generate knockout mutants. Recently, the development of base editors has broadened the scope of genome editing, allowing precise and efficient nucleotide substitutions. In this study, we produced mutants in two cultivated elite cultivars of the tetraploid potato (*Solanum tuberosum*) using stable or transient expression of the CRISPR-Cas9 components to knockout the amylose-producing *StGBSSI* gene. We set up a rapid, highly sensitive and cost-effective screening strategy based on high-resolution melting analysis followed by direct Sanger sequencing and trace chromatogram analysis. Most mutations consisted of small indels, but unwanted insertions of plasmid DNA were also observed. We successfully created tetra-allelic mutants with impaired amylose biosynthesis, confirming the loss-of-function of the StGBSSI protein. The second main objective of this work was to demonstrate the proof of concept of CRISPR-Cas9 base editing in the tetraploid potato by targeting two loci encoding catalytic motifs of the StGBSSI enzyme. Using a cytidine base editor (CBE), we efficiently and precisely induced DNA substitutions in the KTGGL-encoding locus, leading to discrete variation in the amino acid sequence and generating a loss-of-function allele. The successful application of base editing in the tetraploid potato opens up new avenues for genome engineering in this species.

**Key Message:** The *StGBSSI* gene was successfully and precisely edited in the tetraploid potato using gene and base editing strategies, leading to plants with impaired amylose biosynthesis.

## Introduction

Most staple crops are harvested for their starch-storing organs, such as cereal grains, tubers and storage roots. They are mainly cultivated as food or feed for humans or livestock, but there is also an increasing demand of renewable resources for non-food applications (Zeeman et al. 2010). Originating from Latin America, potato (*Solanum tuberosum*) constitutes one of the most important crops for human consumption owing to its starch-rich tubers.

In higher plants, photosynthetic cells produce transitory starch in chloroplasts and also export sucrose to heterotrophic organs, where it is converted to starch for long-term storage in amyloplasts (Lemoine et al. 2013). Starch is an insoluble glucan composed of two polymers of glucose, the ratio of which strongly determines its physiochemical properties. Amylose is a nearly linear glucose polymer with α-1,4-linked residues, whereas amylopectin is made up of α-1,4-linked chains with α-1,6-linkages at branch points, conferring crystallinity to the starch granule. In higher plants, starch biosynthesis is mainly mediated by four classes of enzymes: ADP-glucose pyrophosphorylases, starch synthases (SSs), starch branching enzymes (SBEs) and starch debranching enzymes (SDBEs) (Zeeman et al. 2010). Starch synthases isoforms SSI, SSII, SSIII govern the elongation of the chains of amylopectin. SSIV is involved in starch initiation; the role of SSV and SSVI is still unclear (Helle et al. 2018; Roldan et al. 2007). In addition, granule-bound starch synthase (GBSS) binds to the starch granule and mediates amylose biosynthesis (Ball et al. 1998; Rongine De Fekete et al. 1960).

Amylose determines many physicochemical properties of starch, namely its pasting temperature and viscosity (Bull et al. 2018; Park et al. 2007). Therefore, modifying potato starch composition by decreasing the amylose (or amylopectin) content may be useful for industrial applications. For example, modulating GBSSI function directly *in planta* can lead to the reduction of post-harvest treatments (Sonnewald and Kossmann 2013). In most dicot species, such as potato, the GBSSI protein is encoded by a single nuclear locus (Cheng et al. 2012). This monogenic control has facilitated the production of amylose-free potato varieties through mutational breeding (Hovenkamp-Hermelink et al. 1987; Muth et al. 2008). In some studies, transgenic approaches have been used to silence the *GBSSI* gene through antisense (Kuipers et al. 1994; Visser et al. 1991) or RNAi technologies (Andersson et al. 2003). Although the genetically modified potato Amflora (BASF) has been commercialized for two years, to date the development of such transgenic crops for commercial purposes has been limited in the European Union, mainly due to regulatory policies (Zeeman et al. 2010).

In the past years, genome editing techniques have received much attention due to their powerful applications in model plants and crops. These techniques rely on the precise introduction of DNA double-strand breaks (DSBs) in the plant genome through a variety of techniques (Ma et al. 2016). DSBs are readily recognized by the cell and repaired either through non-homologous end joining (NHEJ) or homologous recombination (HR), the former being the main pathway for the repair of DSBs in somatic cells (Puchta 2005). Contrary to HR, NHEJ is error-prone and may lead to random small insertions or deletions (indels) at the cut site. Since 2012, tremendous breakthroughs have been made using clustered regularly interspaced short palindromic repeats (CRISPR)-associated nuclease Cas9 systems (Sternberg et al. 2016). When expressed in eukaryotic cells, CRISPR-Cas9 systems fulfil their function by forming a complex made up of a single-guide RNA molecule (sgRNA) and the Cas9 nuclease (Jinek et al. 2012). The latter recognizes a protospacer adjacent motif (PAM), mainly NGG in the case of *Streptococcus pyogenes* Cas9 (*Sp*Cas9), which is found just downstream of the target sequence. Recognition of the PAM likely destabilizes the adjacent double-stranded DNA, allowing base pairing between a 17-21 bp target-dependent sequence from the sgRNA and its matching target DNA sequence (Anders et al. 2014; Bortesi and Fischer 2015). The Cas9 nuclease eventually induces DSB in DNA about 3 bp upstream of the PAM sequence. Error-prone NHEJ may lead to small indels, potentially resulting in frameshift mutations (Soyars et al. 2018). A truncated and/or non-functional protein will be translated, possibly triggering the nonsense-mediated decay (NMD) pathway that leads to mRNA degradation (Pauwels et al. 2018; Popp and Maquat 2016).

The CRISPR-Cas9 system has been successfully developed in potato in the past few years (Hameed et al. 2018). For example, the *StIAA2* and the *StALS* genes have been efficiently targeted in a double haploid cultivar and/or a tetraploid potato using *Agrobacterium*-mediated stable transformation (Butler et al. 2015; Butler et al. 2016; Wang et al. 2015). More recently, the full knockout of the *StGBSS* gene in the tetraploid potato cultivar Kuras was obtained using transient expression of CRISPR-Cas9 components in protoplasts, either as DNA plasmids or as ribonucleoprotein (RNP) complexes (Andersson et al. 2017; Andersson et al. 2018).

In addition to generating gene knockout through the introduction of indels, CRISPR-Cas9 can precisely replace nucleotides, allowing the study of specific domains within a protein or also providing polymorphism within a gene to confer valuable agronomic traits in elite varieties. Such modifications can be carried out using HR and the insertion of a DNA template bearing the polymorphism. However, precise and efficient base editing via HR in plants suffers from low efficiency and the delivery of template DNA is still challenging (Schindele et al. 2018). A new CRISPR-Cas9-based genome editing system has recently been developed based on fusing either a cytidine or an adenine deaminase to a Cas9 nickase (nCas9 D10A), leading to a C-to-T or an A-to-G conversion on the edited strand, respectively (Schindele et al. 2018). The sgRNA directs the nCas9/deaminase fusion to the target locus, enabling edition on the non-complementary strand. Cytidine or adenine is converted to uracil or inosine, respectively, without introducing DSBs. Uracil and inosine are then converted through DNA replication to thymine and guanine, respectively, although C-to-G and C-to-A conversions have also been reported (Nishida et al. 2016). The edition has been shown to take place in a 3-8 bp deamination window distal from the PAM on the non-complementary strand, with some variations according to the base editor (Gaudelli et al. 2017; Kang et al. 2018; Shimatani et al. 2017; Zong et al. 2017). The nCas9 nicks the opposite strand of the deamination site to direct DNA repair mechanisms to the G-or-T-containing DNA strand using the edited strand as a template for mismatch repair, thus preserving the edit and increasing the mutation rate (Nishida et al. 2016). To date, base editors have been successfully used in some crops, including rice, tomato, wheat, maize, oilseed rape, potato and watermelon (Hua et al. 2018a; Kang et al. 2018; Li et al. 2018; Shimatani et al. 2017; Tian et al. 2018; Yan et al. 2018; Zong et al. 2018; Zong et al. 2017).

In this work, we targeted the *StGBSSI* gene to assay various CRISPR-Cas9 tools using stable and transient expression in different varieties of the tetraploid potato *Solanum tuberosum*. We identified single- and multi-allelic edited plants, and mutations in all four alleles were observed, resulting in amylose biosynthesis impairment. To induce this modification, we used protoplast transformation, which can produce non-transgenic plants. Finally, we assessed the efficiency of the cytidine base editor (CBE) in two loci encoding catalytic motifs of the StGBSSI enzyme, resulting in precise base conversion and amino-acid substitution. Our results highlight that CRISPR-Cas9 gene and base editing can be efficiently developed in a genetically complex and vegetatively propagated crop such as potato to modify agronomic traits.

## Materials and methods

### Plant material

Potato cultivars Desiree (ZPC, the Netherlands) and Furia (Germicopa, France) were *in vitro* propagated in 1X Murashige and Skoog (MS) medium (pH 5.8) including vitamins (Duchefa, the Netherlands), 0.4 mg/L thiamine hydrochloride (Sigma-Aldrich, USA), 2.5% sucrose and 0.8% agar powder (VWR, USA). Plants were cultivated in a growth chamber at 19°C with a 16:8 h light:dark photoperiod. For the production of *in vitro* microtubers, plants were placed in 5 mL of 1X MS culture medium for about one month at 19°C with a 16:8 h light:dark photoperiod, until depletion of the medium. Then, 4 mL of half-strength MS medium supplemented with 50 μg/mL coumarin and 4 μg/mL kinetin were added and the plants were transferred to a dark room at 20°C for about five additional weeks, until microtubers were sufficiently developed. The production of tubers in soil was carried out by transferring *in vitro* plants to a greenhouse. Watering was stopped 2 weeks before the tubers were harvested.

### Target identification

Genomic sequences of the *StGBSSI* gene (Gene ID from NCBI: 102577459) were obtained from leaf DNA extracted using the NucleoSpin Plant II kit (Macherey-Nagel, Germany) according to the manufacturer’s instructions. Primers were designed using Primer3 (Untergasser et al. 2012) and Netprimer (www.premierbiosoft.com/netprimer) from the reference genome (https://plants.ensembl.org/Solanum_tuberosum/Info/Index), and are listed in Supplementary Table S1. Amplification was carried out on about 10 ng of DNA using Invitrogen Platinum SuperFi DNA polymerase (Thermo Fisher Scientific, USA) following the supplier’s instructions. PCR products were cloned into the pCR4-TOPO TA vector (Thermo Fisher Scientific, USA) and transformed by heat shock into One Shot™ TOP10 Chemically Competent *E. coli* (Thermo Fisher Scientific). Bacteria were grown overnight at 37°C on LB plates with 50 μg/mL kanamycin and plasmids from randomly selected positive clones were purified using QIAprep Spin Miniprep kit (QIAGEN, Germany) and Sanger sequenced (Genoscreen, Lille, France).

Target loci in the *StGBSSI* gene were selected manually based on their distance from the KTGGL and PSRFEPCGL motifs, and then analysed using the CRISPOR software (Haeussler et al. 2016). In this study, four guides were designed upstream of the NGG PAM and were named sgGBSS1 (5’-GTTGGTCCTTGGAGCAAAAC-3’), sgGBSS2 (5’-TTGTCATTACCCGATGTCCG-3’), sgGBSS3 (5’-GGACTAGGTGATGTTCTTGG-3’) and sgGBSS4 (5’-CCAAGCAGATTTGAACCTTG-3’). Guide design was performed according to the predicted efficiency and the off-target potential from the potato reference genome, selecting guides with no off-target site with less than two mismatches and with no mismatch in the seed region adjacent to the PAM.

### Vector construction

For the sgGBSS1 and sgGBSS2 targets, guide sequences were respectively placed downstream a *StU6* (Z17301.1) (Guerineau and Waugh 1993) or a *StU3* (NW_006239017.1) promoter (Supplementary Fig. S1) and upstream a sgRNA scaffold previously described (Shimatani et al. 2017). The constructs were synthesized (Genscript, USA) with Gateway AttB1 and AttB2 sequences on both sides to perform a BP reaction with the pDONR207 plasmid. For stable transformation using *Agrobacterium tumefaciens*, the pDONR207-containing the sgGBSS1 construct was LR-recombined with the pDe-Cas9 (Fauser et al. 2014) harbouring a *nptII* resistance cassette. The resulting plasmid was transferred into *A. tumefaciens* (C58pMP90) strain by heat shock. For transient expression in protoplasts, the pDeCas9, pDONR207-sgGBSS1 and pDONR207-sgGBSS2 were purified using the QIAGEN Plasmid Plus Midi Kit (QIAGEN, Germany), followed by a sodium acetate precipitation.

For the sgGBSS3 and sgGBSS4 targets, guide sequences were cloned into the pDicAID_nCas9-PmCDA_NptII_DELLA (Shimatani et al. 2017). To replace the guide targeting the *SlDELLA* locus, the plasmid was digested by the FastDigest restriction enzymes *Bst*XI and *Spe*I in the presence of Fast Alkaline Phosphatase (Thermo Fischer Scientific, USA). The sgGBSS3 and sgGBSS4 guide sequences were synthesized (Genscript, USA) together with a portion of the AtU6 promoter and the sgRNA scaffold, flanked by a *Bst*XI and a *Spe*I restriction site. This construct was cloned into the pDONR207 plasmid using a BP reaction (Gateway) and digested as described above, without the phosphatase treatment. Guide sequences were then ligated into the digested binary vector using T4 DNA ligase (New England Biolabs, USA). Reaction mixture was transformed into One Shot™ TOP10 Chemically Competent *E. coli* (Thermo Fisher Scientific, USA) and bacteria were grown overnight at 37°C on LB plates containing 100 μg/mL spectinomycin. Plasmids were Sanger sequenced (Genoscreen, Lille, France) and transferred into *A. tumefaciens* C58pMP90 strain by heat shock.

### *Agrobacterium*-mediated transformation

Stem and petiole explants were cut from the top of 3 to 5 week-old *in vitro* plants, and placed overnight in a growth chamber on 1X Murashige and Skoog (MS) medium (pH 5.8) including vitamins (Duchefa, The Netherlands), 2.5% sucrose, 0.4 mg/L thiamine hydrochloride (Sigma-Aldrich, USA), 1 mg/L indole-3-acetic acid (Sigma-Aldrich, USA), 1 mg/L zeatin-riboside (Sigma-Aldrich, USA), 1 mg/L gibberellin A3 (Sigma-Aldrich, USA) and 0.7% agar powder (VWR, USA). *A. tumefaciens* C58pMP90 strain containing CRISPR-Cas9 plasmids was grown overnight at 28°C at 250 rpm in LB medium with 50 μg/mL rifampicin, 25 μg/mL gentamicin and 50 μg/mL spectinomycin. The bacterial optical density (OD) was set to ≈ 0.2 in the MS medium without antibiotics. Potato explants were co-cultured with *A. tumefaciens* for 48 h at 25°C in the dark, and were then washed with sterile water and placed on the culture medium described above, supplemented with 250 μg/mL cefotaxime, 100 μg/mL timentin^®^ and 50 μg/mL kanamycin. After two weeks, explants were transferred onto a fresh culture medium with reduced indole-3-acetic acid (0.1 mg/L), and were then maintained by subculturing every three weeks. Regenerating shoots were transferred to the culture medium containing 50 mg/L kanamycin and/or tested for the presence of the T-DNA by PCR.

### PEG-mediated protoplast transfection

Protoplasts were isolated from leaves of 3-5 week-old plants propagated *in vitro*. Plants were kept in the dark for at least 18 h before the start of the digestion. Protoplast digestion and isolation were mainly performed as previously described by Yoo et al. (2007) with some modifications. Digestion was performed overnight in the dark at 25°C using 0.2% cellulase Onozuka R10 (Yakult Pharmaceutical Industry, Japan) and 0.2% macerozyme R10 (Yakult Pharmaceutical Industry, Japan), without shaking. The next day, the protoplast solution was gently shaken at 70 rpm for 30 min to release round protoplasts. The solution was filtered with a 40 μm cell strainer before the washing steps. Round-bottomed tubes were used during all the procedure. Transient expression of pDeCas9, pDONR207-sgGBSS1 and/or pDONR207-sgGBSS2 was performed by a PEG-mediated transfection using 3.6 × 10^5^ protoplasts in 300 μL. A total amount of 10 μg of plasmid DNA was added to the protoplasts, followed by 300 μL of 25% PEG4000 (Merck, Germany) for 2 min. The transfection solution was gradually diluted in MaMg medium (0.4 M mannitol, 15 mM MgCl_2_, pH 5.8) and kept at 4°C for 15 min in the dark. Protoplasts were embedded in 3 mL of alginate solution (Sigma, USA) as described by Andersson et al. (2017). Plant regeneration was essentially performed following the protocol described by Masson et al. (1987).

### Detection of mutations

For the detection of CRISPR-Cas9-induced deletions and/or insertions, PCR genotyping was performed across sgGBSS1 and sgGBSS2 target sequences. Amplification was carried out using the GoTaq G2 Flexi DNA polymerase (Promega, USA) and the PCR products were run on a 1.5% agarose gel.

The high-resolution melting (HRM) curve analysis was performed as described in Veillet et al. (2016). Genomic DNA from leaf tissue was extracted using the NucleoSpin Plant II kit (Macherey-Nagel) according to the manufacturer’s instructions. Primers were designed to obtain amplicons of about 100 bp (Supplementary Table S1) and tested for their specificity and dimer formation on an agarose gel. PCR amplification was carried out in 12 μL volumes containing 6 μL of High Resolution Melting Master (Roche Applied Science, Germany), 0.24 μL of each 10 μM primer, 1.44 μL of 25 mM MgCl_2_ solution, and 5-30 ng of genomic DNA. PCR was performed using 96-well white PCR plates using the LightCycler^®^ 480 II system (Roche Applied Science, Germany). The amplification started with an initial denaturation step at 95°C for 5 min, followed by 40 cycles of 95°C for 10 s, 63-59°C for 10 s and 72°C for 10 s. HRM was immediately performed with a denaturation step at 95°C for 1 min followed by an incubation at 40°C for 1 min. The melting curve was generated over a 65-95°C range, with a 0.2°C/s increment and 25 acquisitions per °C. Results were analysed with LightCycler^®^ 480 Gene Scanning software (Roche Applied Science, Germany). To detect mutations, all samples were spiked with 10-20% of wild type DNA. For each mutated plant, a new HRM run was carried out with and without spiking to identify putative homozygous mutated plants. All plants with a mutated profile were Sanger sequenced (Genoscreen, Lille, France), as well as some plants with a wild-type profile to check for the sensitivity of the HRM. Some plants harbouring both an HRM profile and a Sanger chromatogram of interest were then analysed by cloning the PCR products into the pCR4-TOPO TA vector (Thermo Fisher Scientific, USA) followed by Sanger sequencing. Using the natural polymorphism found in wild-type Desiree, each sequencing read was paired to a particular allele.

### Inference of CRISPR editing (ICE) analysis

Chromatograms from Sanger sequencing were analysed using the Inference of CRISPR Editing (ICE) software (Synthego, USA), an open-source and free-to-use web-based tool (https://ice.synthego.com) (Hsiau et al. 2018).

### Amylose assay

For a visual assay, 20 μL of a half-strength Lugol’s solution (Sigma-Aldrich, USA) was directly dropped onto the surface of a sliced tuber.

For size-exclusion chromatography analysis of starch polysaccharides, potato starch was isolated and purified according to Helle et al. (2018). Washed and peeled tubers were cut into 0.5 cm x 0.5 cm pieces and ground in a mortar with 10 mL of ultrapure water. Samples were then filtered through a nylon net (100 μm mesh) and starch granules were left to sediment. Starch suspensions were then washed three times prior to size-exclusion chromatography analysis. Starch polysaccharides were separated on a Sepharose CL-2B column as described in Delvalle et al. (2005). Briefly, ≈1-2 mg of starch were dissolved in 500 μL of 10 mM NaOH and loaded on a CL-2B column (0.5 cm x 65 cm). 300 μL fractions were collected at a flow rate of 12 mL/h prior to measuring the OD and λmax of the iodine-polysaccharide complexes with the use of a microplate spectrophotometer.

### Structural analysis of the GBSS mutation

Eight GBSS templates were selected using the Modeller 9.18 programme (Webb and Sali 2016) based on sequence identity (>20%) from *Hordeum vulgare* (PDB:4HLN), *Saccharomyces cerevisiae* (PDB:1YGP), *Oryza sativa* (PDB:3VUE), *Corynebacterium callunae* (PDB:2C4M), *Oryctolagus cuniculus* (PDB:2GJ4), *Streptococcus* (PDB:4L22), *Cyanophora paradoxa* (PDB:6GNG) and *Cyanobacterium sp*. (PDB:6GNF). The best model was chosen using the discrete optimized protein energy (DOPE) method (Shen and Sali 2006) and/or the GA341 method (John 2003; Melo et al. 2002). The model was optimized using energy minimization protocols available in Yasara software (Krieger et al. 2009).

## Results

### Design of CRISPR-Cas9 targets

In potato, the StGBSSI protein is encoded by a single gene that contains 13 exons (Cheng et al. 2012). The loci of interest were sequenced in two cultivars, Desiree and Furia, to assess inter-allelic polymorphism and to design targets (Fig. 1 and Supplementary Fig. S2 and S3). The locus encoding the catalytic domain KTGGL (amino acids 95 to 99), likely involved in ADP-glucose binding (Nazarian-Firouzabadi and Visser 2017), was completely conserved in both cultivars (Fig. 1). Based on these sequences, two different strategies were implemented to target the *StGBSSI* gene using CRISPR-Cas9 editing methods. We designed two sgRNAs in exon 1 and 2, named sgGBSS1 and sgGBSS2 (Fig. 1), to knock out the *StGBSSI* gene using the pDeCas9 construct (Fauser et al. 2014). We also designed one sgRNA in exon 1 (sgGBSS3) and another one in exon 10 (sgGBSS4) to target the loci encoding the KTGGL and the PSRFEPCGL catalytic domains, respectively (Fig. 1), using a CBE. The sgGBSS4 spanned a region of synonymous allelic variation at its 5’ end in one of the four alleles in the Desiree cultivar (Fig. 1).

**Fig. 1.**
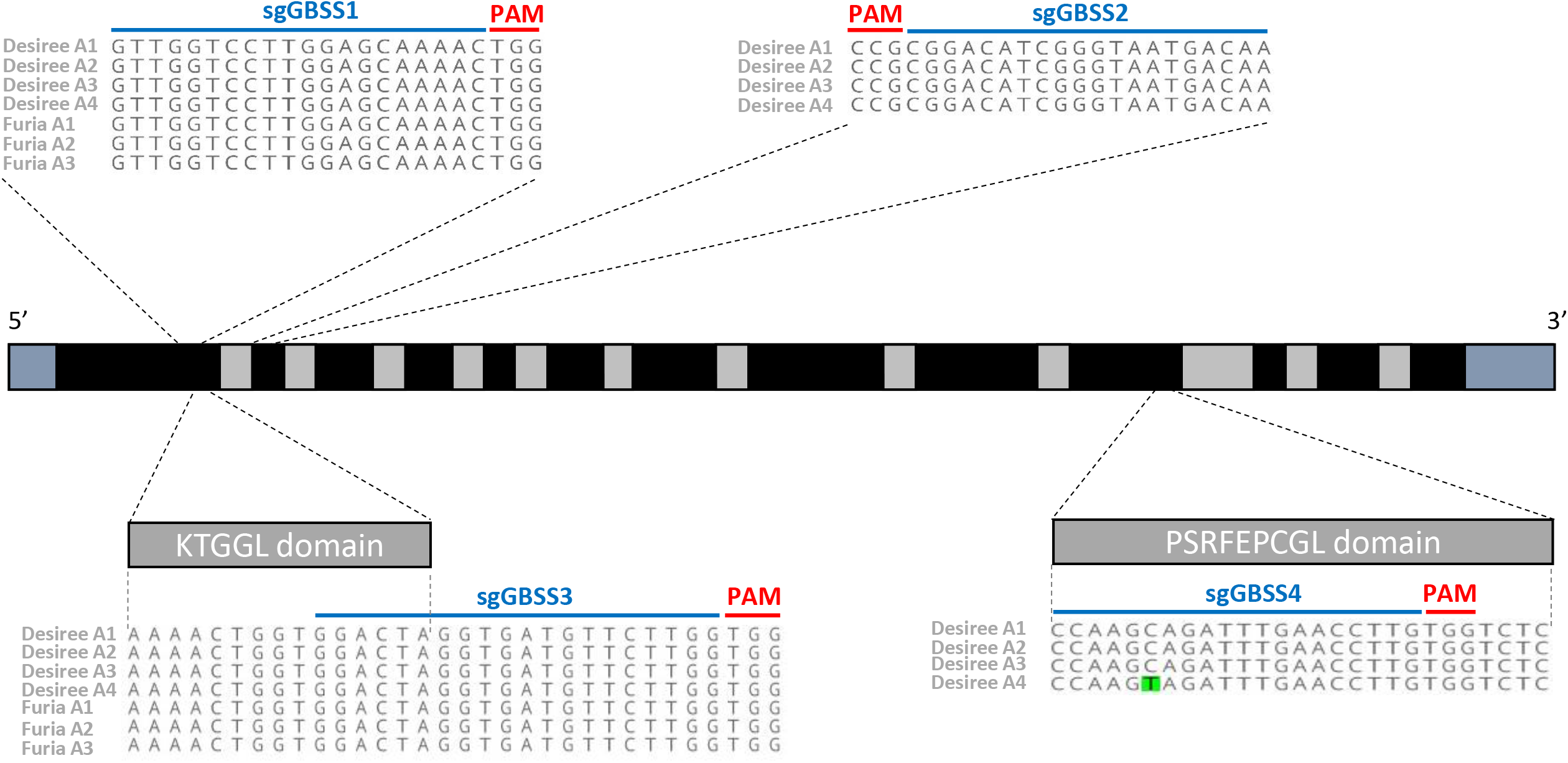
*StGBSSI* structure and CRISPR-Cas9 targets. The *StGBSSI* gene structure is composed of 13 exons (in black) and 12 introns (in grey), with 5’UTR and 3’UTR on both sides (in blue). The localization and sequences of the four targeted loci used in this study are indicated, as well as the allelic variation (in green) in the loci encoding the PSRFEPCGL catalytic domain. The guide sequences (sgGBSS) are depicted in blue and the PAM sequence in red. Polymorphism between allelic variants is not shown.

### Highly efficient *StGBSSI* gene knockout using *Agrobacterium*-based stable transformation

Our objective was to knockout the *StGBSSI* gene through the creation of small indels at the locus encoding the KTGGL domain, targeted by sgGBSS1, following *Agrobacterium*-mediated transformation. After transformation of Desiree explants, 21 kanamycin-resistant transgenic plantlets were regenerated. Mutations were screened using HRM analysis: 15 out of 21 Desiree transgenic plantlets showed a distinct melting-curve shape, resulting in a 71% editing efficiency at the sgGBSS1 target (Fig. 2a). Because the frequency of mosaic plants is often high in primary transformants (Fauser et al. 2014; Pan et al. 2016; Peng et al. 2017), we cloned PCR amplicons before sequencing to obtain individual sequences. Two primary transformants were selected, for which 18 independent sequences were analysed. In both plants, small deletions were detected in the *StGBSS* targeted site a few bp upstream of the PAM (Fig. 2b). In some cases, deletions downstream of the PAM were also observed, as previously reported in potato (Butler et al. 2015; Wang et al. 2015). Mutations were characterised for four and three alleles for the 17T.701.008 and 17T.701.010 plants, respectively. Interestingly, several different sequences (up to six) were observed for a single allele from the same plant, demonstrating that these primary transformants were mosaic (Fig. 2b). Depending on the chimerism level, non-mosaic mutants may be obtained in clonally propagated progenies. In all cases, the mutation was predicted to induce a frameshift or to alter the KTGGL domain, likely leading to a loss of function of the encoded protein.

**Fig. 2.**
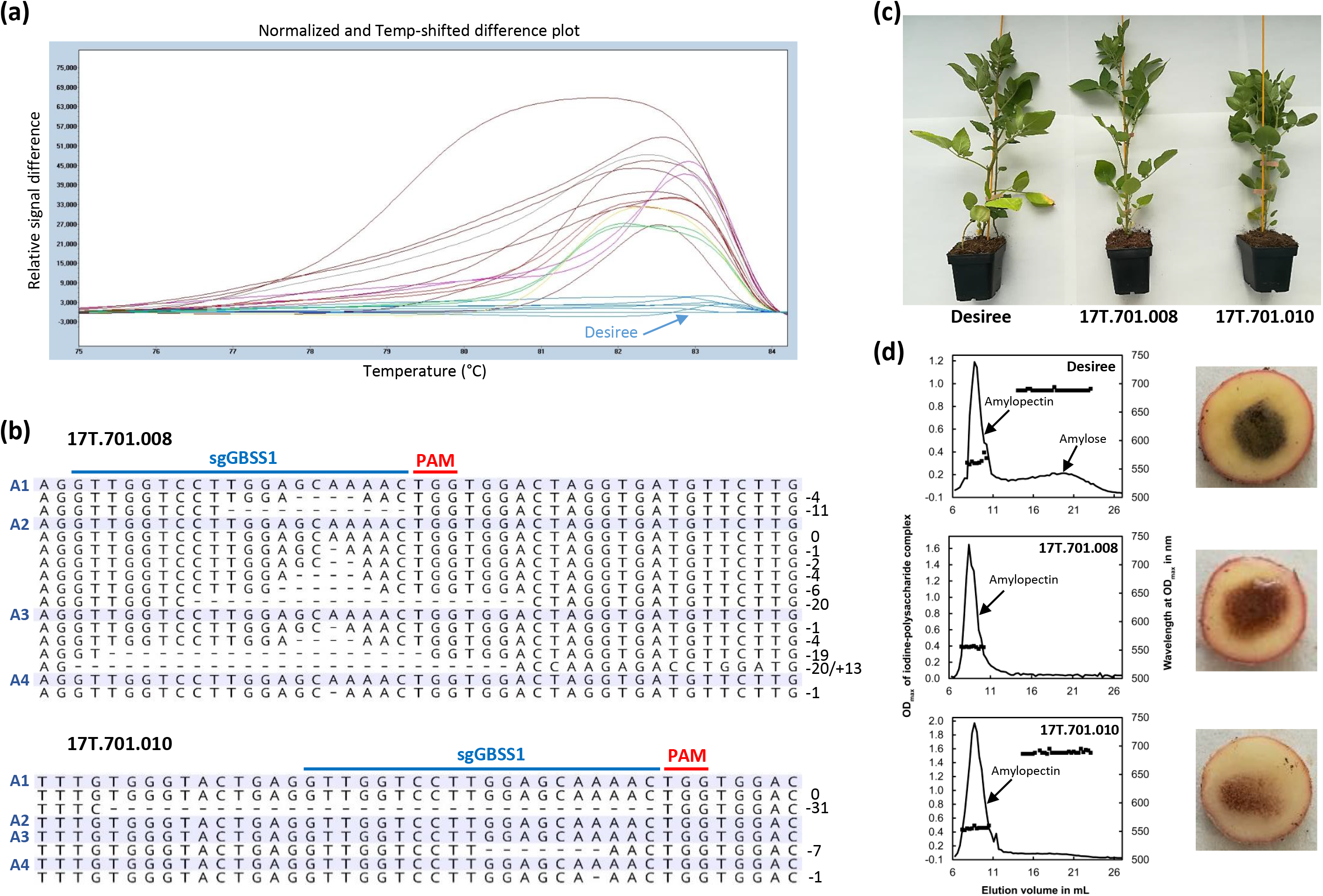
Generation and identification of CRISPR-Cas9-mediated mutations in primary transformants of potato cultivar Desiree. **(a)** High-resolution melting (HRM) analysis of *Agrobacterium*-transformed potato plants. Desiree sample was defined as the base curve (blue line). Non-mutated plants are shown in blue and the mutated ones are shown in different colours, according to the shape of their melting curve. **(b)** Alignment of sequencing reads from *StGBSSI* target locus of two independent mutated primary transformants with the sequences from their respective wild-type alleles (named A1/2/3/4, and highlighted in blue). Natural polymorphism between the four Desiree allelic variants is outside of the represented window. The length of deletion (−) or insertion (+) is indicated on the right of each read. The PAM motif is shown in red and the sgGBSS1 target locus in blue. **(c)** Picture of about 2 month-old potato plants grown in soil in a greenhouse with a natural photoperiod. **(d)** Determination of amylose content in tubers harvested from soil-grown potato with a rapid iodine test (pictures) and with size-exclusion chromatography (graph). Amylopectin and amylose peaks are indicated with black arrows.

To explore the phenotypic consequences of the targeted mutations, the two mutated plants were transferred to a greenhouse with a natural photoperiod to produce tubers in soil. We did not notice any obvious deleterious effects of the mutation to overall plant growth (Fig. 2c). Tubers were harvested after three months and assessed for their amylose content. First, the iodine solution was directly dropped onto a tuber slice for a quick qualitative analysis, staining brown for the two mutated plants but dark-blue for wild-type plants (Fig. 2d), as expected for starch containing amylose. Starch polysaccharides were then separated by size-exclusion chromatography using a Sepharose CL-2B matrix. Although the amylopectin peak (high mass fraction) was unaffected in the two mutants compared to Desiree, amylose accumulation (low mass fraction) was totally abolished or strongly impaired in the 17T.701.008 and 17T.701.010 plants, respectively (Fig. 2d). The residual amylose content in 17T.701.010 may result from the presence of a sufficient amount of cells with unedited alleles (Fig. 2b). Taken together, these data clearly show that *StGBSSI* can be efficiently knocked-out in the tetraploid potato by CRISPR-Cas9, leading to modifications in tuber starch quality.

### Successful gene editing in regenerated potato plants using protoplast transfection

*Agrobacterium*-based gene editing is associated with the integration of the *Cas9*-harbouring transgene. To generate transgene-free edited plants through transient expression of CRISPR components, Desiree protoplasts were transfected using plasmid DNA. In this case, we simultaneously applied two sgRNAs (sgGBSS1 and sgGBSS2) on opposite DNA strands and spaced 135 bp apart, aiming at generating larger deletions. Four months after transfection, 444 plantlets were regenerated and their DNA was extracted from leaf samples.

Despite the use of the two sgRNAs, only 2 regenerated plants out of 269 (0.7%) showed a PCR band shift consistent with the expected deletion for at least one allele (Fig. 3a). This result indicates that simultaneous cutting by both guides is low. However, 6 out of 269 plants (2.2%) were detected with a clear insertion in the targeted region (Fig. 3a). We also analysed the 17T.716.146 plant, which displayed no wild-type alleles, but both smaller and larger band shifts in the targeted locus. The locus was cloned and sequenced, revealing the expected deletion of 142 bp as well as two insertions of 116 bp and 211 bp, originating from the plasmids used in the experiment (data not shown). This is similar to previous observations made in potato (Andersson et al. 2017). In line with the absence of the wild-type *StGBSSI* allele, the amylopectin content was not affected in soil-grown tubers from 17T.716.146 plants (no growth penalties observed), and amylose accumulation was completely abolished (Fig. 3b). This result confirms the loss of function of the *StGBSSI* gene in 17T.716.146 plants.

**Fig. 3.**
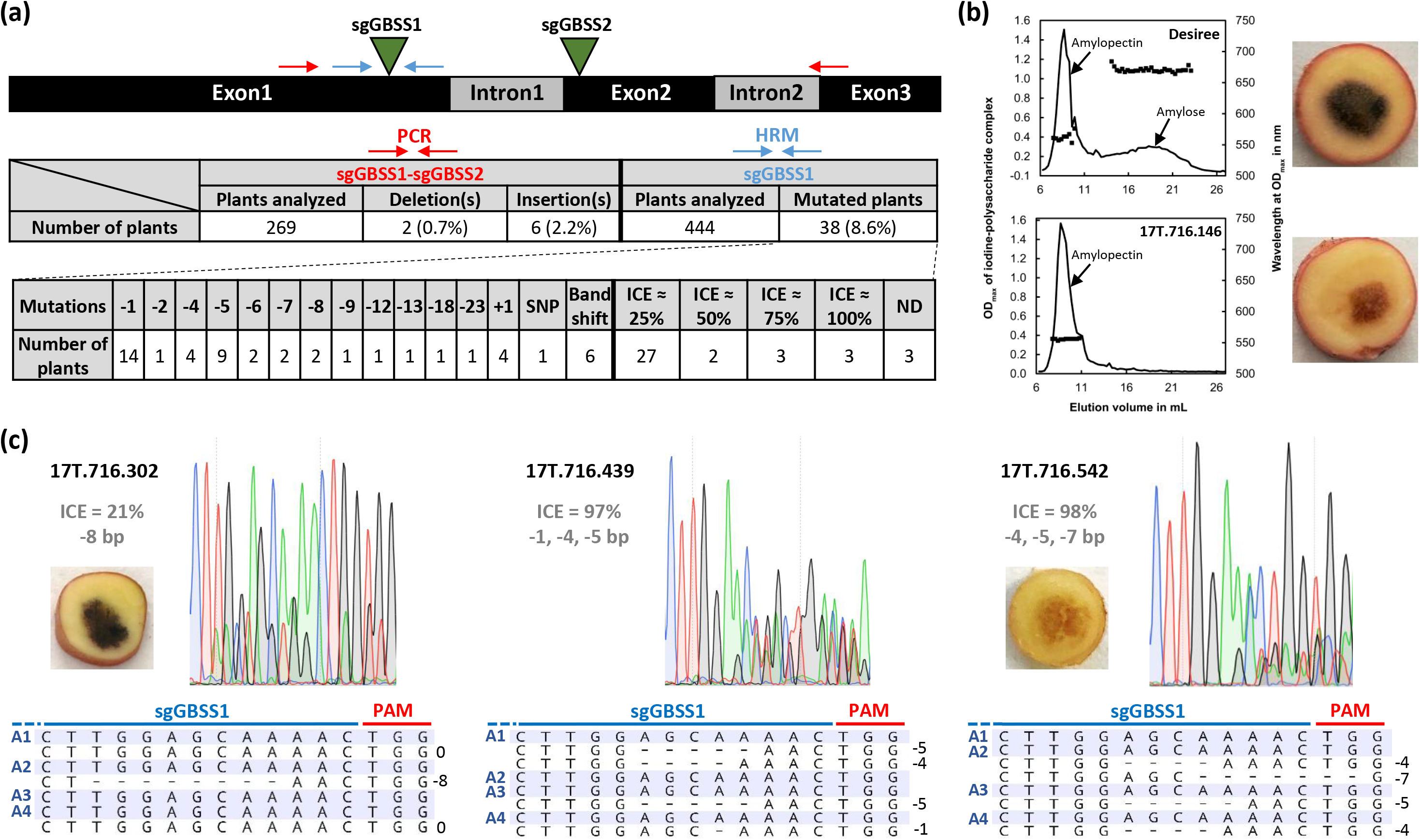
Generation and identification of CRISPR-Cas9-mediated *StGBSSI* mutations in regenerated plants from PEG-mediated transfection of potato cultivar Desiree protoplasts. **(a)** Genotyping strategies for selecting deletions in *StGBSSI*. The PCR band shift assay using primers on both sides of the targeted region (sgGBSS1 and sgGBSS2) is shown in red, and the HRM analysis at the sgGBSS1-targeted site is given in blue. Exons and introns are shown in dark and grey, respectively. The frequency of mutated plants was calculated based on the number of regenerated and analysed plants. The nature of mutations was determined after Sanger sequencing using both manual analysis and ICE software. Most plants could be assigned an ICE score (≈ 25, 50, 75 or 100%). **(b)** Determination of amylose content in tubers harvested from soil-grown potato plants using a rapid iodine test (pictures) and with size-exclusion chromatography (graph). Amylopectin and amylose peaks are indicated with black arrows. **(c)** Sanger sequencing chromatograms (C in blue, G in dark, A in green and T in red) and Sanger sequencing reads from a few regenerated plants at the sgGBSS1-targeted locus. The ICE score and the mutation are indicated in grey. The wild-type allelic sequences are highlighted in blue (A1/2/3/4). Polymorphism between Desiree allelic variants is not located within the sequencing window. The length of deletion (−) is indicated on the right of each read. The PAM motif is shown in red and the sgGBSS1-targeted locus in blue. Determination of amylose content with a rapid iodine test in tubers harvested from soil-grown (17T.716.302) or from *in vitro*-propagated (17T.716.542) potatoes is shown on the left of the chromatogram.

Due to the low efficiency of inducing large deletions using the two-guide strategy, we first used HRM analysis to detect small indels at the sgGBSS1-cutting site: 38 plants out of 444 (8.6%) were identified as differing from Desiree at the *StGBSSI* locus (Fig. 3a). For these 38 plants, the expected polymorphism was assessed by simultaneously sequencing the four loci by direct Sanger sequencing. The resulting sequences were analysed with ICE software, which determines the rate of CRISPR-Cas9 editing at a specific locus, allowing us to reconstruct the mutated alleles. Most of the mutations consisted of small indels (−23 to +1 bp) leading to frameshifts or amino-acid deletions (Fig. 3a). The region targeted by sgGBSS2 could not be similarly analysed because the high natural polymorphism downstream the target prevented us from designing suitable HRM primers (Supplementary Fig. S4).

In a tetraploid species, one single allele is sufficient to produce amylose (Andersson et al. 2017). Using ICE software, the mutants can be theoretically classified in four categories in a tetraploid species like potato: ICE 25% (one allele likely to be mutated), ICE 50% (2 alleles likely mutated), ICE 75% (3 alleles likely mutated) and ICE 100% (4 alleles likely mutated). We therefore screened the 38 *StGBSSI* mutant plants to identify plants with all four alleles mutated. As a control, we cloned and sequenced 10 individual amplicons from the 17.716.302 plant, which displayed a 21% ICE score: in agreement with this score, only one allele was modified, associated with a 8 bp deletion. Accordingly, tubers from this plant were not affected in amylose accumulation (Fig. 3c). Most of the confirmed edited plants were similarly mutated in a single allele (29 plants, 77%), but we focused on three plants (9%) mutated at all four alleles (Fig. 3a and Supplementary Fig. S5), including the aforementioned 17T.716.146 plant, identified as knocked-out. For the two putative tetra-allelic mutants (17T.716.439 and 17T.716.542), we also cloned and sequenced the sgGBSS1 target locus. No wild-type sequence could be detected among the 20 sequencing reads obtained for each plant. The mutations identified were in accordance with those predicted by the ICE software, with 5, 4 and 1 bp deletions for 17T.716.439, and 7, 5 and 4 bp deletions for 17T.716.542 (Fig. 3c and Supplementary Fig. S5). We detected two different mutations for the same allele in 17T.716.439 and 17T.716.542, indicating that these mutants may be mosaic (Fig. 3c). Surprisingly, in both plants, sequences corresponding to only three different natural allelic variants could be detected, and the A2 and A1 alleles were not detected in 17T.716.439 and 17T.716.542, respectively (Fig. 3c). One possible explanation may be a very large insert in the cutting site of one allele, preventing PCR detection. Alternatively, a change in chromosome number/structure due to somaclonal variation, which is common during plant regeneration from protoplasts (Fossi et al. 2019), potentially explaining the stunted growth of these two plants. The rapid amylose assay performed on an *in vitro* microtuber from 17T.716.542 indicates that this mutant was strongly impaired in amylose biosynthesis (Fig. 3c).

To assess the effectiveness of the protoplast strategy, we then wanted to ensure that no foreign DNA was inserted elsewhere in the plant genome. We performed PCR on the mutated plants with four couples of primers matching the CRISPR/Cas9 plasmids. Interestingly, although we did not observe any amplification for most of the plants (32 plants, 84%), four of them (11%) had integrated at least one large plasmid fragment (>350 bp) (Supplementary Fig. S6). This was confirmed by the sequencing of some of the amplicons. Two of the plants that harboured the *nptII* amplicon successfully grew on a medium containing kanamycin, confirming the presence of a functional *nptII* resistance cassette (Supplementary Fig. S7). We postulate that at least some of these insertions may result from random integration into the genome, although we cannot exclude the insertion of a very large fragment into the target site (preventing PCR amplification) or into an off-target site.

To summarize, the two-guide strategy did not improve the efficiency of mutagenesis, but we were able to edit and knockout the *StGBBSI* gene in regenerated plantlets by transiently expressing CRISPR-Cas9 components into potato protoplasts. In comparison with the *Agrobacterium*-mediated knockout approach, we were able to regenerate plants without large insertions of T-DNA fragments, although foreign DNA insertions could be detected in a subset of plants. However, smaller insertions may be present as part of the substantial chromosomic reshuffling caused by protoplast regeneration(Fossi et al. 2019). Finally, a high number of regenerated plants displayed stunted growth (20 to 40%), confirming the major drawback of protoplast-based regeneration.

As a direct application of our work on Desiree, a cultivar commonly used in laboratories for many years, we targeted the *StGBSSI* gene through protoplast transfection in the Furia cultivar, a recently registered starch potato. We applied a single-guide approach with sgGBSS1 only and, four months after transfection, we detected 42 out of 259 regenerated plants (16%) as mutated at this locus using HRM (Fig. 4a). In total, eight plants (19% of the mutants) were identified by Sanger sequencing as potential tetra-allelic mutants (Fig. 4b, c). This suggests that our strategy can be readily developed in other cultivars for plant molecular breeding, but with the same potential drawbacks of somaclonal variation and residual foreign DNA integration (data not shown).

**Fig. 4.**
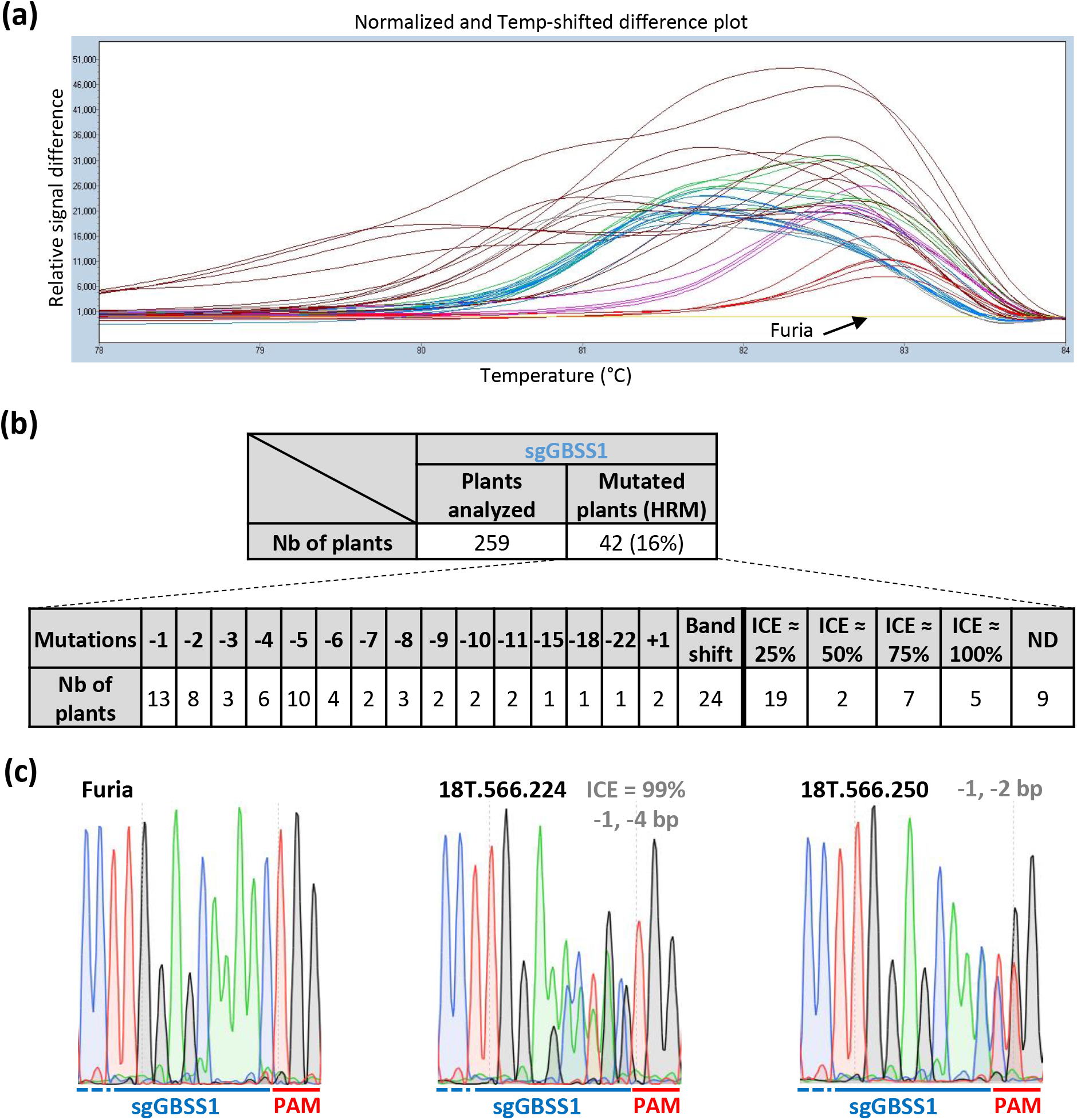
Generation and identification of CRISPR-Cas9-mediated *StGBSSI* mutations in regenerated plants from PEG-mediated transfection of potato cultivar Furia protoplasts. **(a)** High-resolution melting (HRM) analysis of regenerated potato plants. Furia sample was defined as the base curve (yellow). Mutated plants are shown in different colours, according to the shape of their melting curve. **(b)** Summary of the mutation detection strategy. Frequency of mutated plants was calculated based on the number of regenerated and analysed plants. The nature of mutations was determined after Sanger sequencing using a manual analysis and ICE software. Most plants could be assigned an ICE score (≈ 25, 50, 75 or 100%). **(c)** Sanger sequencing chromatograms from Furia and two mutated plants at the sgGBSS1-targeted locus (C in blue, G in dark, A in green and T in red). The ICE score and the mutation are indicated in grey for each callus. The PAM motif is indicated in red and the sgGBSS1-targeted locus in blue.

### Precise and efficient base editing using a CRISPR-Cas9 cytidine deaminase fusion

Recently, substitution of nucleotides without DSBs has been successfully carried out on plants using base editing tools. Here, we used a cytidine base editor (Shimatani et al. 2017) to target two loci encoding catalytic domains of the StGBSSI enzyme. One suitable guide sequence in each target locus was designed, named sgGBSS3 (targeting the KTGGL encoding locus) and sgGBSS4 (targeting the PSRFEPCGL encoding locus) (Fig. 1). We carried out two independent *Agrobacterium*-mediated explant transformations in Desiree, leading to the regeneration of 48 and 15 transgenic plants targeting sgGBSS3 and sgGBSS4 sequences, respectively. HRM analysis suggests that both sgGBSS3- and sgGBSS4-targeted sites were efficiently mutated with 43 out of 48 transgenic plants and 13 out of 15 transgenic plants edited, respectively. Direct Sanger sequencing confirmed these results, demonstrating a very high mutation efficiency, close to 90% (Fig. 5a).

**Fig. 5.**
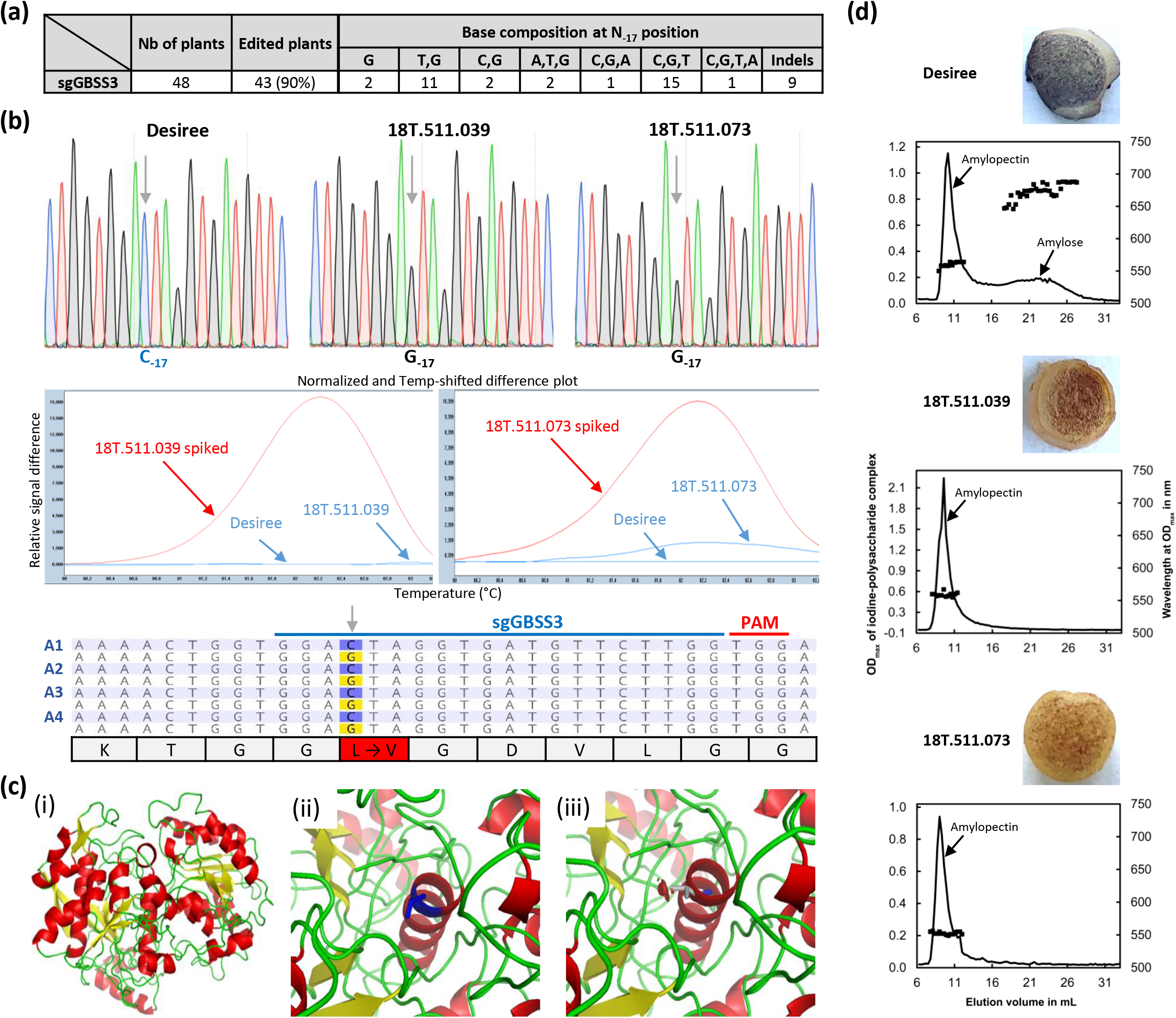
Generation and identification of CBE-mediated mutations in primary transformants of potato cultivar Desiree. **(a)** Base-editing efficiencies at sgGBSS3 target in primary transformants. The number of plants harbouring a specific nucleotide conversion is indicated for C_−17_. **(b)** Sanger sequencing results and high-resolution melting (HRM) outputs for two edited plants at the sgGBSS3 target harbouring a perfect C_−17_-to-G_−17_ substitution. The HRM analysis was performed without or with the addition of a small amount of Desiree DNA in the samples (spiked). Non-mutated profiles are shown in blue and the mutated ones are shown in red, according to the shape of the melting curve. The chromatograms are the result of direct sequencing of all four alleles together. Grey arrows indicate the location of base substitutions. Sanger reads were aligned with the wild-type allelic sequences (in blue). Natural polymorphism between Desiree allelic variants is not shown. Base substitutions are highlighted in yellow. The PAM motif is indicated in red and the sgGBSS3-targeted locus in blue. Amino-acid residues for all four alleles of the edited plants are indicated below the Sanger reads. Changes in amino-acid residues are highlighted in red. K: Lysine, T: Threonine, G: Glycine, L: Leucine, V: Valine, D: Aspartic acid. **(c)** Localisation of mutated residue in the 3D model of StGBSSI. Overall 3D model of StGBSSI (i), zoom on L99 (ii) and the L99V mutation (iii). The StGBSSI consists of 24 α-helix (in red) and 19 β-strands (in yellow). L99 is shown in blue (ii). L99 (situated in α-helix5) is mutated in V99 and the clash between the side chain from V99 and other residues is indicated by a red square (iii). **(d)** Determination of amylose content in microtubers harvested from *in vitro* plants with a rapid iodine test (picture) and with a size-exclusion chromatography (graph). Amylopectin and amylose peaks are indicated with black arrows.

The sgGBSS4-targeted site had been selected to assess the edition efficiency of three closely located Cs in the edition window (C_−20_, C_−19_ and C_−15_, counting from the PAM). Sanger sequencing showed that these 3 Cs can be substituted with a T, and to a lesser extent with a G or an A, but strikingly indels were very frequently found (77%), mainly in the edition window (data not shown). Moreover, we confirmed that, consistent with the presence of a mismatch with one allele at the distal end of the guide (Fig. 1), complete editing of the four alleles could not be characterized in the sgGBSS4 target zone.

For the sgGBSS3-targeted site, Sanger sequencing revealed that only C_−17_ in the editing window was most frequently substituted with a G or a T, and to a much lesser extent with an A (Fig 5a-b). In all edited plants, we observed a C_−17_-to-G_−17_ substitution, while a C_−1_7-to-T_−17_ transition occurred in 30 mutated plants (70%) (Fig. 5a and Supplementary Fig. S8). C_−17_-to-A_−17_ conversion was detected in only four plants (9%). Among the 43 mutants, indels were observed in nine plants (21%), with an initiation site mainly located in the vicinity of the C_−17_ (Supplementary Fig. S8). As expected, we did not observe base substitution at positions surrounding the edition window. In particular, we identified a perfect C_−17_-to-G_−17_ conversion in two plants (18T.511.039 and 18T.511.073) (Fig. 5a-b). Interestingly, the complete homozygous conversion of the target base hinders detection by HRM, a fact that can be circumvented by the addition of a small amount of wild-type DNA into the samples (spiking) to clearly alter the melting shape (Fig. 5b). The cloning and sequencing of the targeted region in both plants confirmed that these plants were tetra-allelic for the mutation and non-mosaic (Fig. 5b). This C_−17_-to-G_−17_ base substitution leads to the replacement of the leucine (L) by a valine (V) in the KTGGL motif, potentially leading to a loss of function of the protein. The StGBSSI 3D protein structure was modelled based on available 3D structures (see Methods) and the mutated residue L99V was reported on the GBSSI 3D model (Fig 5c). It is located within an α-helix and is likely to impair its structure, subsequently leading to abnormal GBSSI protein folding and/or impacting the glucose binding. The amylose assay on *in vitro* microtubers supports this assumption, because amylose accumulation was totally impaired in the two base-edited plants compared with wild-type Desiree (Fig. 5d), indicating that the leucine residue in the KTGGL is essential for StGBSSI catalytic activity.

## Discussion

### Gene editing using CRISPR-Cas9 is efficient in elite cultivars of tetraploid potato

Our work demonstrates that the delivery of CRISPR-Cas9 components either through stably integrated transgene(s) or transient expression of plasmids can efficiently induce targeted mutations into the *StGBSSI* gene in elite cultivars of the tetraploid potato. Using *Agrobacterium*-mediated transformation, we successfully obtained, in the first generation, tetra-allelic mutants with impaired amylose biosynthesis, confirming the loss of function of the StGBSSI enzyme (Fig. 2). For fundamental research purposes, the persistence of the transgene(s) may not be a problem because Tang et al. (2018) showed that the continued presence of CRISPR-Cas9 reagents in rice does not cause off-target mutations if sgRNAs are rigorously designed.

In sexually propagated species, the stably integrated transgene can be eliminated through Mendelian segregation, generating transgene-free lines (Ricroch et al. 2017). However, the segregation of the transgene is not feasible in the vegetatively propagated cultivated potato, because selfing would change the cultivar characteristics of this highly heterozygous species. Thus, regenerated mutants lacking unwanted insertion of foreign DNA are of utmost interest, especially for commercial applications. With this goal, we used the transient expression of CRISPR-Cas9 plasmids to generate transgene-free knockout mutants. While we conducted these experiments, two studies were published showing how the *StGBSSI* gene can be knocked-out in plantlets regenerated from protoplasts of the Kuras cultivar, using both plasmid DNA and RNPs, respectively (Andersson et al. 2017; Andersson et al. 2018). Our results are consistent with these findings, extending them to other cultivars and using different transfection and screening strategies. Although being less efficient than *Agrobacterium*-mediated stable transformation, we demonstrated that PEG-mediated transformation of Desiree and Furia protoplasts is efficient for the generation of tetra-allelic edited plants at a reasonable rate (8 to 19% of all mutated plants), producing plants impaired in amylose biosynthesis.

At the same time, our work highlights caveats associated with genome editing approaches in the highly heterozygous tetraploid potato. Rigorous analysis of the target locus must be carried out to avoid polymorphism that will impair tetra-allelic editing (as shown here for sgGBSS4) or genotyping (as exemplified here by the locus downstream from sgGBSS2). Similarly, we confirm here that a dual sgRNA approach on opposite DNA strands does not necessarily lead to an efficient deletion rate of a large segment of the gene, probably because the cutting efficiencies at the two sgRNA-targeted sites are not similar. For an efficient and cost-effective screening of mutants, we present a streamlined approach with an HRM analysis followed by direct Sanger sequencing of positive plants and accompanied by sequence analysis mediated by the ICE algorithm.

During the protoplast transfection process, the delivery of CRISPR-Cas9 components through plasmids may result in the insertion of degraded DNA fragments into the target sequence (Salomon and Puchta 1998), but also into random sites (Kim et al. 2017; Liang et al. 2017). Accordingly, we found that a substantial rate of mutants harboured insertions in the target sequence, which may originate from plasmid DNA and/or host genome. These results corroborate previous studies using transient transfection in potato protoplasts, revealing a high rate of DNA insertions (Andersson et al. 2018; Clasen et al. 2016). Moreover, the unpredictable insert sequence can impair the detection of foreign DNA insertions into the plant genome, likely underestimating the rate of random and unwanted insertions. Whole-genome sequencing of the mutated line(s) may be an exhaustive - and expensive - option, as done in tomato (Nekrasov et al. (2017) and in rice (Tang et al. (2018). Another strategy for CRISPR-Cas9 expression relies on the use of CRISPR-Cas9 ribonucleoproteins (RNP). This method has been successfully developed in some plant species, including potato, completely avoiding the risk of foreign DNA insertion(s) into the host genome (Andersson et al. 2018; Liang et al. 2017; Woo et al. 2015). Recently, Chen et al. (2018) developed a method using an *Agrobacterium*-mediated transient CRISPR-Cas9 gene expression system to generate transgene-free mutants without the need for sexual segregation, which holds great potential for vegetatively propagated crops. Furthermore, we developed a strategy for the production of T-DNA free edited plants using the *Agrobacterium*-mediated delivery of a CBE, opening up new perspectives for genome engineering, especially in vegetatively propagated species like potato (Veillet et al. 2019). The use of nanomaterials for CRISPR components delivery also holds great promises for genome editing without foreign DNA integration (Demirer et al. 2019). These promising strategies, which directly generate transgene-free edited plants without the need for transgene segregation, may provide science-based evidences for decision-makers and may also help to reduce public concerns about gene edited crops.

Somaclonal variation in regenerated plantlets from protoplasts or *Agrobacterium* transformation constitutes another bottleneck for an efficient gene editing strategy, especially for commercial purposes. For example, this type of variation can result in changes in the number of chromosomes from polyploidy to aneuploidy, chromosome rearrangements and DNA base deletions and substitutions (Krishna et al. 2016). In our conditions, we observed a substantial rate of plants with stunted growth or abnormal development, which may be due to somaclonal/epigenetic variation (Fossi et al. 2019). Collectively, our results suggest that many regenerated plants need to be screened to isolate clean edited plants suitable for commercial purposes. Therefore, the improvement of the transformation/regeneration method is crucial to obtain a higher ratio of tetra-allelic mutants.

### CRISPR-Cas9 cytidine deaminase fusion precisely and efficiently edits targeted bases

Base editing using CBE has been recently developed in some plant species, allowing the precise modification of cytidine residues in a small edition window. In this study, we successfully applied CBE to the tetraploid potato using *Agrobacterium*-mediated transformation. We obtained a high rate of edited plants for the two targeted-sites, close to 90% of the transgenic plants. By targeting the active site KTGGL, we achieved a perfect C_−17_-to-G_−17_ substitution in all the alleles of two independent plants, leading to a L99V substitution in this motif. This amino acid change, which is predicted to alter the function of StGBSSI, effectively led to amylose biosynthesis impairment. Although this mutation resulted in a loss-of-function allele, such precise genome editing approach is very likely to help create new allelic variants with potentially modified activity. This is a promising result for the characterization of specific protein motifs in potato, but also for crop improvements through the production of gain-of-function mutants, for example exhibiting resistance to plant pathogens at no yield loss (Bastet et al. 2019). Based on our results, we suggest that further studies are needed to draw up the guidelines for designing more efficient sequence guides. One such avenue to explore is whether C-rich regions are likely to be associated with indels, which should be avoided for precise gene editing. Similarly, instead of the expected C-to-T mutation, C-to-G/-A conversion may be attractive for the diversity of amino-acid substitutions, but unwanted in some cases. To avoid such undesired products due to uracil excision and downstream repair systems, the addition of a uracil DNA glycosylase inhibitor protein (UGI) to the construct is a promising approach, because this strategy has been shown to result in a majority of C-to-T conversions and a reduced rate of indel formation (Nishida et al. 2016). Furthermore, the recent application of base editing in potato using a fusion of human APOBEC3A, nCas9 and UGI demonstrated that C-to-T conversion is efficiently mediated in a 17 bp edition window, independently of the sequence context and with a very low frequency of indels or undesired edits (Zong et al. 2018).

Base editing technology will benefit from the expanding base editing toolbox. In particular, an adenine base editor (ABE) has been developed, mediating clean A-to-G substitutions (Schindele et al. 2018). Together with CBEs, this new ABE opens up further possibilities for fine-tuning allele variation. Nevertheless, a recent study pointed out that CBEs, but not ABEs, can generate genome-wide off-target mutations (Jin et al. 2019), highlighting the need for an optimization of CBEs and/or the use of delivery methods reducing the expression of CRISPR components to a few days. Finally, CRISPR-based base editing requires the presence of a PAM sequence that places the target base(s) within a small base-editing window. This requirement limits the number of sites that can be targeted in plant genomes, as experienced in our work for the loci encoding catalytic domains of the StGBSSI. To overcome this limitation, the commonly used SpCas9 nickase can be replaced by Cas9 orthologues from other bacterial and archeal species that display alternative PAM compatibilities. For example, *SpCas9* orthologues from *Streptococcus thermophilus* and *Staphylococcus aureus* have been successfully used for gene editing in *Arabidopsis* (Steinert et al. 2015). Furthermore, (Hua et al. 2018b) developed new CBEs and ABEs with engineered SpCas9 and SaCas9 nickase variants that considerably expand the targetable sites in the rice genome. Great efforts have been made recently in engineering expanded PAM SpCas9 variants (xCas9 and Cas9-NG) with broadened PAM compatibility in mammalian and plant cells (Endo et al. 2018; Hu et al. 2018; Nishimasu et al. 2018; Wang et al. 2018), opening new exciting avenues for base editing.

### Conclusion

Our present work confirms that the *StGBSSI* gene can be successfully targeted and altered in elite potato cultivars using gene and base editing systems. Therefore, the *StGBSSI* appears to be a very good - and economically feasible - gene model to assess genome editing in potato, as well as its potential future technological improvements. Furthermore, these results can be transferred to other starch-producing crops: the targeted mutagenesis of *GBSS* was recently shown to be efficient in modifying amylose content in cassava (Bull et al. 2018). With the extraordinarily rapid development of genome editing tools in both animals and plants, researchers are now able to efficiently characterize genes underlying important agronomic traits directly into crops, at the base-pair level. Along with the development of plant transformation and regeneration processes, there is no doubt that genome editing will have a tremendous impact on basic research and on molecular crop improvement.

## Author contributions

FV, LC and JEC designed the experiments. FV, LC, MPK, ZT, FS, MM and NS performed the experiments. FV and JLG wrote the article. FV, LC, FS, NS, PD, JLG and JEC discussed the data and revised the article. All authors approved the final manuscript.

## Funding

This work was supported by the INRA UMR IGEPP and the Investissement d’Avenir program of the French National Agency of Research for the project GENIUS (ANR-11-BTBR-0001_GENIUS). ZT is funded by a CIFRE PhD grant from SYNGENTA.

## Conflict of interest statement

The authors declare that the research was conducted in the absence of any commercial or financial relationships that could be construed as a potential conflict of interest.

## Acknowledgments

We thank Dr Puchta and his team (Botanical Institute II, Karlsruhe Institute of Technology, Karlsruhe, Germany) for providing the pDeCas9 plasmid via Marianne Mazier (INRA-UR1052, Montfavet, France) and to Keiji Nishida for providing the pDicAID_nCas9-PmCDA_NptII_Della plasmid. We thank Fabien Nogué and Peter Rogowsky for their efficient management of the GENIUS project and their constructive discussions. We are grateful to Gabriel Guihard for his contribution in the sequencing of Desiree alleles. We thank the BrACySol BRC (INRA Ploudaniel, France) that provided us with the plants that were used in this study and Gisèle Joly and Catherine Chatot from Florimond Desprez/Germicopa (France) for helpful discussions and choice of the starch elite cultivar Furia. We are grateful to Jean-Louis Rolot from CRAW (Belgium) for providing us with the procedure to obtain *in vitro* microtubers as well as to Marie-Ange Dantec and all the INRA Ploudaniel greenhouse staff for acclimation and cultivation of the regenerated plantlets. Finally, we are thankful to Carolyn Engel-Gautier and Marina Perez Benitez for their help in correcting the manuscript.

