## Supplemental materials for "The *Solanum tuberosum GBSSI* gene: a target for assessing gene and base editing in tetraploid potato"

### Supplementary data

**Supplementary Fig. S1: Sequences of StU6 and StU3 promoters used in this study.**

**StU6 promoter:**

GTTCAGTTGCATTATGTCTTTATACACCCCTACCTAGATGATTAAGTTTTACTTTA
GTTGGTGTGAAATGGATAAATTCTAAAATATGAGGTGTGGAATGAAGGATTGTCA
TCAATTAGTTGGCCCCAACCAAGTAAAATAAGAAGGCCGGCCCATTACAATTAAG
TCGTCACACAAGTGGGCTTCATTGAAACAAGCGCAAAAAGGAGTCCAGGCCCGT
GTTAGCGTGAAGACTCAACCAGCGATTTCTCCCTCATCGGTTGCACAGAAAAGCT
GTGTGTTGTTTATATGGCGAAACCTAACAGTCTGACTTG

**StU3 promoter:**

TTCTTTAATGACTCGGAATATGAAATTTCCAATTTTTATAATTTTTGATTGGTTTAT
ATCTTTTTTTTAAATTTTTTCTTTCCTAATTTATTATGGAGTTCATGGCAAGAGGAA
AAACTGCCTACTGGGAATTGGAACGGTTTCTACGAAATTAACTATCTACACGTTA
AAATAAAATTAATGCATAATTATTATTTTTTCTATAACAAATCAAAAACTGAAAT
CTGACATAAATTTTATTACTTTTATTGCACTTTAGCCTTAGAGATTGCGTAGTAGC
CAGCGTAGCTGTGTCAGGGGCCAATAGATTGTTCCCACATCGGCAGTGCAACACA
TAAGCTCCAGCTTTATGAGGCTTCTTACCAGGAGCATATGTCA

**Supplementary Fig. S2: Allelic variants for the beginning of the *StGBSSI* gene in Desiree and Furia**

**Desiree allelic variant 1:**

TCATCTATTCTGATTTTGATTCTCTTGCCTACTGTAATCGGTAATAAATGTGAATGCTTCCT
CTTCTTCTTCTTCTCAGAAATCAATTTCTGTTTTGTTTTTGTTCATCTGTAGCTTATTCTCTG
GTAGATTCCCCTTTTTGTAGACCACACATCACATGGCAAGCATCACAGCTTCACACCACTT
TGTGTCAAGAAGCCAAACTTCACTAGACACCAAATCAACCTTGTCACAGATAGGACTCAG
GAACCATACTCTGACTCACAATGGTTTAAGGGCTGTTAACAAGCTTGATGGGCTCCAATC
AACAACTAATACTAAGGTAACACCCAAGATGGCATCCAGAACTGAGACCAAGAGACCTG
GATGCTCAGCTACCATTGTTTGTGGAAAGGGAATGAACTTGATCTTTGTGGGTACTGAGG
TTGGTCCTTGGAGCAAAACTGGTGGACTAGGTGATGTTCTTGGTGGACTACCACCAGCCC
TTGCAGTAAGTCTTTCTTTCATTTGGTTACCTACTCATTCATTACTTATTTTGTTTAGTTAG
TTTCTACTGCATCAGTCTTTTTATCATTTAGGCCCGCGGACATCGGGTAATGACAATATCC
CCCCGTTATGACCAATACAAAGATGCTTGGGATACTGGCGTTGCGGTTGAGGTACATCTT
CCTATATTGATACGGTACAATATTGTTCTCTTACATTTCCTGGTTCAAGAATGTGATCATC
TG

**Desiree allelic variant 2:**

TCATCTATTCTGATTTTGATTCTCTTGCCTACTGAATTTGACCCTACTGTAATCGGTGATAA
ATGTGAATGCTTCCTCTTCTTCTTCTTCTTCTCAGAAATCAATTTCTGTTTTGTTTTTGTTCA
TCTGTAGCTTGGTAGATTCCCCTTTTTGTAGACCACACATCACATGGCAAGCATCACAGCT
TCACACCACTTTGTGTCAAGAAGCCAAACTTCACTAGACACCAAATCAACCTTGTCACAG
ATAGGACTCAGGAACCATACTCTGACTCACAATGGTTTAAGGGCTGTTAACAAGCTTGAT
GGGCTCCAATCAAGAACTAATACTAAGGTAACACCCAAGATGGCATCCAGAACTGAGAC
CAAGAGACCTGGATGCTCAGCTACCATTGTTTGTGGAAAGGGAATGAACTTGATCTTTGT
GGGTACTGAGGTTGGTCCTTGGAGCAAAACTGGTGGACTAGGTGATGTTCTTGGTGGACT
ACCACCAGCCCTTGCAGTAAGTCTTTCATTTGGTTACCTACTCATTCATTACTTATTTTGTT
TAGTTAGGTTCTACTGCATCAGTCTTTTTATCATTTAGGCCCGCGGACATCGGGTAATGAC
AATATCCCCCCGTTATGACCAATACAAAGATACTTGGGATACTAGCGTTGCGGTTGAGGT
ACATCTTTCTATATTGATACGGTACAATATTGTTCTCTTACATTTCCTGATTCAAGAATGT
GATCCGCTACTTTATCTG

**Desiree allelic variant 3:**

TCATCTATTCTGATTTTGATTCTCTTGCCTACTGTAATCGGTGATAAATGTGAATGCTTCCT
CTTCTTCTTCTTCTTCTCAGAAATCAATTTCTGTTTTGTTTTTGTTCATCTGTAGCTTGGTAG
ATTCCCCTTTTTGTAGACCACACATCACATGGCAAGCATCACAGCTTCACACCACTTTGTG
TCAAGAAGCCAAACTTCACTAGACACCAAATCAACCTTGTCACAGATAGGACTCAGGAA
CCATACTCTGACTCACAATGGTTTAAGGGCTGTTAACAAGCTTGATGGGCTCCAATCAAG
AACTAATACTAAGGTAACACCCAAGATGGCATCCAGAACTGAGACCAAGAGACCTGGAT
GCTCAGCTACCATTGTTTGTGGAAAGGGAATGAACTTGATCTTTGTGGGTACTGAGGTTG
GTCCTTGGAGCAAAACTGGTGGACTAGGTGATGTTCTTGGTGGACTACCACCAGCCCTTG
CAGTAAGTCTTTCATTTGGTTACCTACTCATTCATTACTTATTTTGTTTAGTTAGTTTCTAC
TGCATCAGTCTTTTTATCATTTAGGCCCGCGGACATCGGGTAATGACAATATCCCCCCGTT
ATGACCAATACAAAGATGCTTGGGATACTAGCGTAGCGGTTGAGGTACATCTTCCTATAT

TGATACGGTACAATATTGTTCCCTTACATTTCCTGATTCAAGAATGTGATCCGCTACTTTA
TCTG

**Desiree allelic variant 4:**

TCATCTATTCTGATTTTGATTCTCTTGCCTACTGAATTTGACCCTACTGTAATCGGTGATAA
ATGTGAATGCTTCTTCTTCTTCTCAGAAATCAATTTCTGTTTTGTTTTTGTTCATCTGTAGC
TTATTCTCTGGTAGATTCCCCTTTTTGTAGACCACACATCACATGGCAAGCATCACAGCTT
CACACTTTGTGTCAAGAAGCCAAACTTCACTAGACACCAAATCAACCTTGTCACAGATAG
GACTCAGGAACCATACTCTGACTCACAATGGGTTAAGGGCTGTTAACAAGCTTGATGGGC
TCCAATCAACAACTAATACTAAGGTAACACCCAAGATGGCATCCAGAACTGAGACCAAG
AGACCTGGATGCTCAGCTACCATTGTTTGTGGAAAGGGAATGAACTTGATCTTTGTGGGT
ACTGAGGTTGGTCCTTGGAGCAAAACTGGTGGACTAGGTGATGTTCTTGGTGGACTACCA
CCAGCCCTTGCAGTAAGTCTTTCATTTGGTTACCTACTCATTCATTACTTATTTTGTTTAGT
TAGTTTCTACTGCATCAGTCTTTTTATCATTTAGGCCCGCGGACATCGGGTAATGACAATA
TCCCCCCGTTATGACCAATATAAAGATGCTTGGGATACTAGCGTTGCGGTTGAGGTACTC
CTTCCTATATTGGTACAACAATATTGTTCTCTTCCATTTCCTGATTCAAGAATGTGATCCG
CTACTTTATCTG

**Furia allelic variant 1:**

TCATCTATTCTGATTTTGATTCTCTTGCCTACTGTAATCGGTGATAAATGTGAATGCTTCCT
CTTCTTCTTCTTCTTCTCAGAAATCAATTTCTGTTTTGTTTTTGTTCATCTGTAGCTTGGTAG
ATTCCCCTTTTTGTAGACCACACATCACATGGCAAGCATCACAGCTTCACACCACTTTGTG
TCAAGAAGCCAAACTTCACTAGACACCAAATCAACCTTGTCACAGATAGGACTCAGGAA
CCATACTCTGACTCACAATGGTTTAAGGGCTGTTAACAAGCTTGATGGGCTCCAATCAAG
AACTAATACTAAGGTAACACCCAAGATGGCATCCAGAACTGAGACCAAGAGACCTGGAT
GCTCAGCTACCATTGTTTGTGGAAAGGGAATGAACTTGATCTTTGTGGGTACTGAGGTTG
GTCCTTGGAGCAAAACTGGTGGACTAGGTGATGTTCTTGGTGGACTA

**Furia allelic variant 2:**

TCATCTATTCTGATTTTGATTCTCTTGCCTACTGTAATCGGTAATAAATGTGAATGCTTCCT
CTTCTTCTTCTTCTCAGAAATCAATTTCTGTTTTGTTTTTGTTCATCTGTAGCTTATTCTCTG
GTAGATTCCCCTTTTTGTAGACCACACATCACATGGCAAGCATCACAGCTTCACACCACTT
TGTGTCAAGAAGCCAAACTTCACTAGACACCAAATCAACCTTGTCACAGATAGGACTCAG
GAACCATACTCTGACTCACAATGGTTTAAGGGCTGTTAACAAGCTTGATGGGCTCCAATC
AACAACTAATACTAAGGTAACACCCAAGATGGCATCCAGAACTGAGACCAAGAGACCTG
GATGCTCAGCTACCATTGTTTGTGGAAAGGGAATGAACTTGATCTTTGTGGGTACTGAGG
TTGGTCCTTGGAGCAAAACTGGTGGACTAGGTGATGTTCTTGGTGGACTA

**Furia allelic variant 3:**

TCATCTATTCTGATTTTGATTCTCTTGCCTACTGTAATCGGTGATAAATGTGAATGCTTCCT
TTCTTCTCAGAAATCAATTTCTGTTTTGTTTTTGTTCATCTGTAGCTTATTCTCTGGTAGAT
TCCCCTTTTTGTAGACCACACATCACATGGCAAGCATCACAGCTTCACACCACTTTGTGTC
AAGAAGCCAAACTTCACTAGACACCAAATCAACCTTGTCACAGATAGGACTCAGGAACC
ATACTCTGACTCACAATGGTTTAAGGGCTGTTAACAAGCTTGATGGGCTCCAATCAACAA

CTAATACTAAGGTAACACCCAAGATGGCATCCAGAACTGAGACCAAGAGACCTGGATGC
TCAGCTACCATTGTTTGTGGAAAGGGAATGAACTTGATCTTTGTGGGTACTGAGGTTGGT
CCTTGGAGCAAAACTGGTGGACTAGGTGATGTTCTTGGTGGACTA

**Supplementary Fig. S3: Allelic variants for exon 10, intron 10 and exon 11 of the *StGBSSI* gene in Desiree**

**Desiree allelic variant 1:**

AAGCTAAAGGAGTGGCAAAATTCAATGTCCCTTTGGCTCACATGATCACTGCTGGTGCTG
ATTTTATGTTGGTTCCAAGCAGATTTGAACCTTGTGGTCTCATTCAGTTACATGCTATGCG
ATATGGAACAGTAAGAACCATAAGAGCTTGTACCTTTTTACTGAGTTTTAAAAAAAGAAT
CACAAGACCTTGTTTTCCGTCTAAAGGTTACTAGCCAACTAAATGTTACTGCAGCAAGCTT
TTCATTTCTGAAAATTGGTTATTTGATTTTAACATAATCACATGTGAGTCAGGTGCCAATC
TGTGCATCGACTGGTGGACTTGTTGACACTGTGAAAGAAGGCTATACTGGATTC

**Desiree allelic variant 2:**

AAGCTAAAGGAGTGGCAAAATTCAATGTCCCTTTGGCTCACATGATCACTGCTGGTGCTG
ATTTTATGTTGGTTCCAAGCAGATTTGAACCTTGTGGTCTCATTCAGTTACATGCTATGCG
ATATGGAACAGTAAGAACCATAAGAGCTTGTACCTTTTTACTGAGTTTTAAAAAAAGAAT
CATAAGACCTTGTTTTCCGTCTAAAGTTTAATAGCCAACTAAATGTTACTGCAGCAAGCTT
TTCATTTCTGAAAATTGGTTATCTAATTTTAACATAATCACATGTGAGTCAGGTGCCAATC
TGTGCATCGACTGGTGGACTTGTTGACACTGTGAAAGAAGGCTATACTGGATTC

**Desiree allelic variant 3:**

AAGCTAAAGGAGTGGCAAAATTCAATGTCCCTTTGGCTCACATGATCACTGCTGGTGCTG
ATTTTATGTTGGTTCCAAGCAGATTTGAACCTTGTGGTCTCATTCAGTTACATGCTATGCG
ATATGGAACAGTAAGAACCATAAGAGCTTGTACCTTTTTACTGAGTTTTAAAAAAAGAAT
CATAAGACCTTGTTTTCCATCTAAAGTTTAACAGCCAACTAAATGTTACTGCAGCAAGCTT
TTCATTTCTGAAAATTGGTTATCTGATTTTAACATAATCACATGTGAGTCAGGTGCCAATC
TGTGCATCGACTGGTGGACTTGTTGACACTGTGAAAGAAGGCTATACTGGATTC

**Desiree allelic variant 4:**

AAGCTAAAGGAGTGGCAAAATTCAATGTCCCTTTGGCTCACATGATCACTGCTGGTGCTG
ATTTTATGTTGGTTCCAAGTAGATTTGAACCTTGTGGTCTCATTCAGTTACATGCTATGCG
ATATGGAACAGTAAGAACCATAAGAGCTTGTACCTTTTTACTGATTTTTAAAAAAAGAAT
CATAAGACCTTGTTTTCCGTCTAAAGTTTAATAGCCAACTAAATGTTACTGCAGCAAGCTT
TTCATTTCTGAAAATTGGTTATCTGATTTTAACATAATCACATGTGAGTCAGGTGCCAATC
TGTGCATCGACTGGTGGACTTGTTGACACTGTGAAAGAAGGCTATACTGGATTC

**Supplementary Fig. S4: Schematic view of the locus downstream of the sgGBSS2-targeted site in Desiree.**

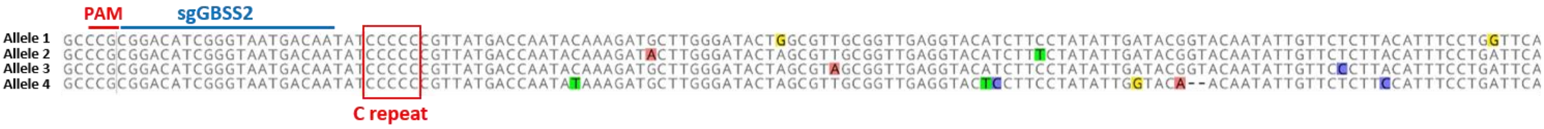

**Supplementary Fig. S5: ICE analysis outputs for CRISPR-Cas9 edited plants in Desiree.** After detection of mutants by HRM analysis, a direct Sanger sequencing was performed. Chromatograms were then analysed using the ICE software (Synthego, USA). Outputs examples for some plants are depicted. Wild-type sequence is indicated by a yellow cross.

## 17T.716.724

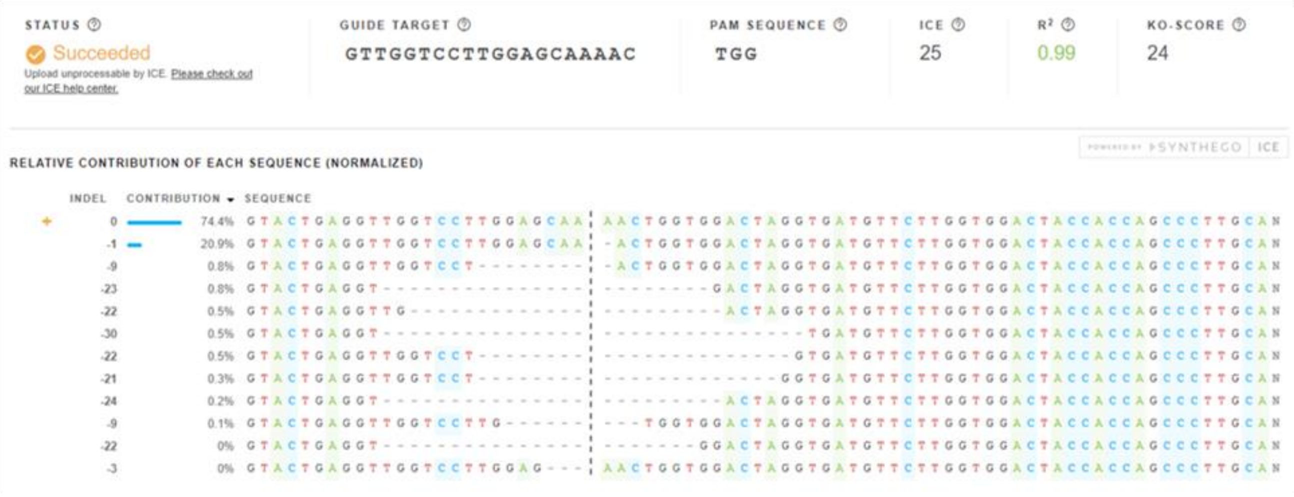


## 17T.716.302

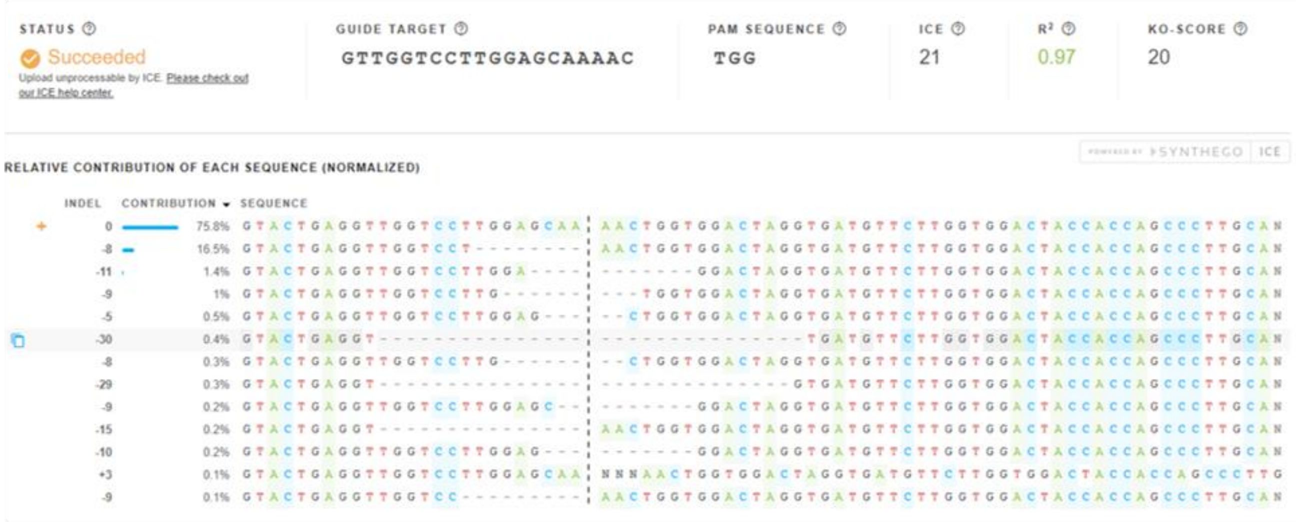


## 17T.716.301

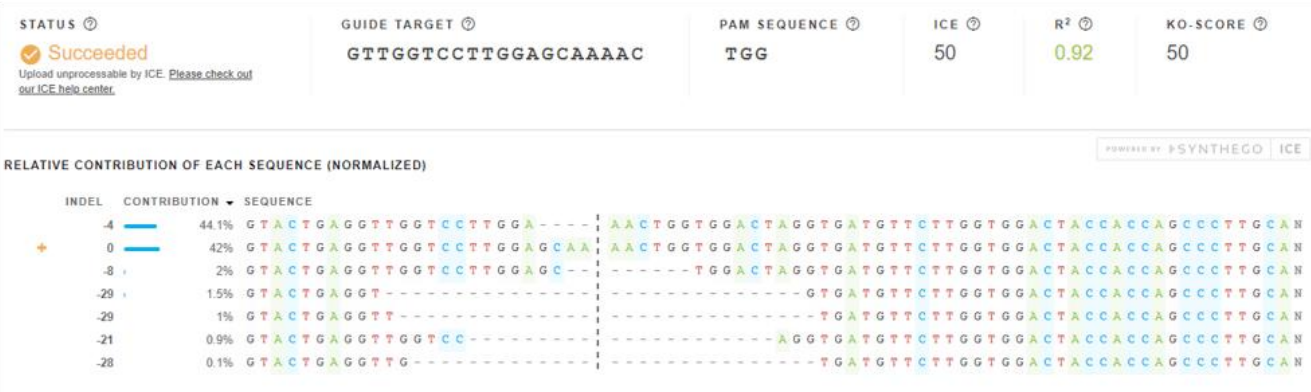

## 17T.716.488

| STATUS | GUIDE TARGET | PAM SEQUENCE | ICE | $R^2$ | KO-SCORE |
|---|---|---|---|---|---|
| Succeeded<br>Upload unprocessable by ICE. Please check out our ICE help center. | GTTGGTCCTTGGAGCAAAAC | TGG | 76 | 0.96 | 72 |

POWERED BY SYNTHEGO ICE

RELATIVE CONTRIBUTION OF EACH SEQUENCE (NORMALIZED)

| INDEL | CONTRIBUTION | SEQUENCE |
|---|---|---|
| -5 | 68.7% | GTACTGAGGTTGGTCCTTGGA---- ¦ -ACTGGTGGACTAGGTGATGTTCTTGGTGGACTACCACCAGCCCTTGCAN |
| 0 | 19.9% | GTACTGAGGTTGGTCCTTGGAGCAA ¦ AACTGGTGGACTAGGTGATGTTCTTGGTGGACTACCACCAGCCCTTGCAN |
| -6 | 3.3% | GTACTGAGGTTGGTCCTTGGAGCA- ¦ -----GTGGACTAGGTGATGTTCTTGGTGGACTACCACCAGCCCTTGCAN |
| -5 | 2% | GTACTGAGGTTGGTCCTTGGAGCA- ¦ ----GGTGGACTAGGTGATGTTCTTGGTGGACTACCACCAGCCCTTGCAN |
| -22 | 0.9% | GTACTGAGGT--------------- ¦ -------GGACTAGGTGATGTTCTTGGTGGACTACCACCAGCCCTTGCAN |
| +12 | 0.7% | GTACTGAGGTTGGTCCTTGGAGCAA ¦ NNNNNNNNNNNNAACTGGTGGACTAGGTGATGTTCTTGGTGGACTACCAC |
| -7 | 0.4% | GTACTGAGGTTGGTCCTTG------ ¦ -ACTGGTGGACTAGGTGATGTTCTTGGTGGACTACCACCAGCCCTTGCAN |
| -7 | 0% | GTACTGAGGTTGGTCCTTGGAGCA- ¦ ------TGGACTAGGTGATGTTCTTGGTGGACTACCACCAGCCCTTGCAN |

## 17T.716.562

| STATUS | GUIDE TARGET | PAM SEQUENCE | ICE | $R^2$ | KO-SCORE |
|---|---|---|---|---|---|
| Succeeded<br>Upload unprocessable by ICE. Please check out our ICE help center. | GTTGGTCCTTGGAGCAAAAC | TGG | 73 | 0.97 | 51 |

POWERED BY SYNTHEGO ICE

RELATIVE CONTRIBUTION OF EACH SEQUENCE (NORMALIZED)

| INDEL | CONTRIBUTION | SEQUENCE |
|---|---|---|
| -1 | 26.5% | GTACTGAGGTTGGTCCTTGGAGCA- ¦ AACTGGTGGACTAGGTGATGTTCTTGGTGGACTACCACCAGCCCTTGCAN |
| 0 | 23.8% | GTACTGAGGTTGGTCCTTGGAGCAA ¦ AACTGGTGGACTAGGTGATGTTCTTGGTGGACTACCACCAGCCCTTGCAN |
| +1 | 22.7% | GTACTGAGGTTGGTCCTTGGAGCAA ¦ NAACTGGTGGACTAGGTGATGTTCTTGGTGGACTACCACCAGCCCTTGCA |
| -9 | 17% | GTACTGAGGTTGGTCCTTGGA---- ¦ -----GTGGACTAGGTGATGTTCTTGGTGGACTACCACCAGCCCTTGCAN |
| -9 | 5.1% | GTACTGAGGTTGGTCC--------- ¦ AACTGGTGGACTAGGTGATGTTCTTGGTGGACTACCACCAGCCCTTGCAN |
| -23 | 1% | GTACTGAGGT--------------- ¦ --------GACTAGGTGATGTTCTTGGTGGACTACCACCAGCCCTTGCAN |
| -4 | 0.4% | GTACTGAGGTTGGTCCTTGGAG--- ¦ -ACTGGTGGACTAGGTGATGTTCTTGGTGGACTACCACCAGCCCTTGCAN |
| -21 | 0.1% | GTACTGAGGTTG------------- ¦ --------GACTAGGTGATGTTCTTGGTGGACTACCACCAGCCCTTGCAN |
| -30 | 0% | GTACTGAGGT--------------- ¦ ---------------TGATGTTCTTGGTGGACTACCACCAGCCCTTGCAN |

## 17T.716.439

| STATUS | GUIDE TARGET | PAM SEQUENCE | ICE | $R^2$ | KO-SCORE |
|---|---|---|---|---|---|
| Succeeded<br>Upload unprocessable by ICE. Please check out our ICE help center. | GTTGGTCCTTGGAGCAAAAC | TGG | 97 | 0.97 | 96 |

POWERED BY SYNTHEGO ICE

RELATIVE CONTRIBUTION OF EACH SEQUENCE (NORMALIZED)

| INDEL | CONTRIBUTION | SEQUENCE |
|---|---|---|
| -5 | 35.6% | GTACTGAGGTTGGTCCTTGG----- ¦ AACTGGTGGACTAGGTGATGTTCTTGGTGGACTACCACCAGCCCTTGCAN |
| -4 | 33.1% | GTACTGAGGTTGGTCCTTGGA---- ¦ AACTGGTGGACTAGGTGATGTTCTTGGTGGACTACCACCAGCCCTTGCAN |
| -1 | 25.9% | GTACTGAGGTTGGTCCTTGGAGCAA ¦ -ACTGGTGGACTAGGTGATGTTCTTGGTGGACTACCACCAGCCCTTGCAN |
| -27 | 0.8% | GTACTGAGGT--------------- ¦ ------------AGGTGATGTTCTTGGTGGACTACCACCAGCCCTTGCAN |
| +2 | 0.5% | GTACTGAGGTTGGTCCTTGGAGCAA ¦ NNAACTGGTGGACTAGGTGATGTTCTTGGTGGACTACCACCAGCCCTTGC |
| -29 | 0.3% | GTACTGAGGT--------------- ¦ --------------GTGATGTTCTTGGTGGACTACCACCAGCCCTTGCAN |
| 0 | 0.3% | GTACTGAGGTTGGTCCTTGGAGCAA ¦ AACTGGTGGACTAGGTGATGTTCTTGGTGGACTACCACCAGCCCTTGCAN |
| +12 | 0.2% | GTACTGAGGTTGGTCCTTGGAGCAA ¦ NNNNNNNNNNNNAACTGGTGGACTAGGTGATGTTCTTGGTGGACTACCAC |
| -23 | 0.1% | GTACTGAGGT--------------- ¦ --------GACTAGGTGATGTTCTTGGTGGACTACCACCAGCCCTTGCAN |
| -22 | 0.1% | GTACTGAGGTTGGTCC--------- ¦ -------------GGTGATGTTCTTGGTGGACTACCACCAGCCCTTGCAN |
| -3 | 0% | GTACTGAGGTTGGTCCTTGGAGC-- ¦ -ACTGGTGGACTAGGTGATGTTCTTGGTGGACTACCACCAGCCCTTGCAN |
| -22 | 0% | GTACTGAGGT--------------- ¦ -------GGACTAGGTGATGTTCTTGGTGGACTACCACCAGCCCTTGCAN |

## 17T.716.542

| STATUS | GUIDE TARGET | PAM SEQUENCE | ICE | $R^2$ | KO-SCORE |
|---|---|---|---|---|---|
| Succeeded<br>Upload unprocessable by ICE. Please check out our ICE help center. | GTTGGTCCTTGGAGCAAAAC | TGG | 98 | 0.98 | 98 |

POWERED BY SYNTHEGO ICE

RELATIVE CONTRIBUTION OF EACH SEQUENCE (NORMALIZED)

| INDEL | CONTRIBUTION | SEQUENCE |
|---|---|---|
| -4 | 53.3% | GTACTGAGGTTGGTCCTTGGA---- ¦ AACTGGTGGACTAGGTGATGTTCTTGGTGGACTACCACCAGCCCTTGCAN |
| -5 | 18.5% | GTACTGAGGTTGGTCCTTGGA---- ¦ -ACTGGTGGACTAGGTGATGTTCTTGGTGGACTACCACCAGCCCTTGCAN |
| +1 | 8.3% | GTACTGAGGTTGGTCCTTGGAGCAA ¦ NAACTGGTGGACTAGGTGATGTTCTTGGTGGACTACCACCAGCCCTTGCA |
| -7 | 7.2% | GTACTGAGGTTGGTCCTTGGAGC-- ¦ -----GTGGACTAGGTGATGTTCTTGGTGGACTACCACCAGCCCTTGCAN |
| -7 | 4.1% | GTACTGAGGTTGGTCCTTGGAGCAA ¦ -------GGACTAGGTGATGTTCTTGGTGGACTACCACCAGCCCTTGCAN |
| -7 | 3.2% | GTACTGAGGTTGGTCCTTGGAGCA- ¦ ------TGGACTAGGTGATGTTCTTGGTGGACTACCACCAGCCCTTGCAN |
| -4 | 2.7% | GTACTGAGGTTGGTCCTTGGAGC-- ¦ --CTGGTGGACTAGGTGATGTTCTTGGTGGACTACCACCAGCCCTTGCAN |
| +13 | 0.4% | GTACTGAGGTTGGTCCTTGGAGCAA ¦ NNNNNNNNNNNNNAACTGGTGGACTAGGTGATGTTCTTGGTGGACTACCA |
| -25 | 0.3% | GTACTGAGGT--------------- ¦ ----------CTAGGTGATGTTCTTGGTGGACTACCACCAGCCCTTGCAN |
| -7 | 0.3% | GTACTGAGGTTGGTCCTTGG----- ¦ --CTGGTGGACTAGGTGATGTTCTTGGTGGACTACCACCAGCCCTTGCAN |

**Supplementary Fig. S6: PCR genotyping for the presence of plasmid DNA in CRISPR-Cas9 edited plants (Desiree).** Only lines of interest have been shown on these agarose gels. Positive control (control +: 17T.701.008 mutant stably transformed with *Agrobacterium*) and negative controls (Desiree) have been included.

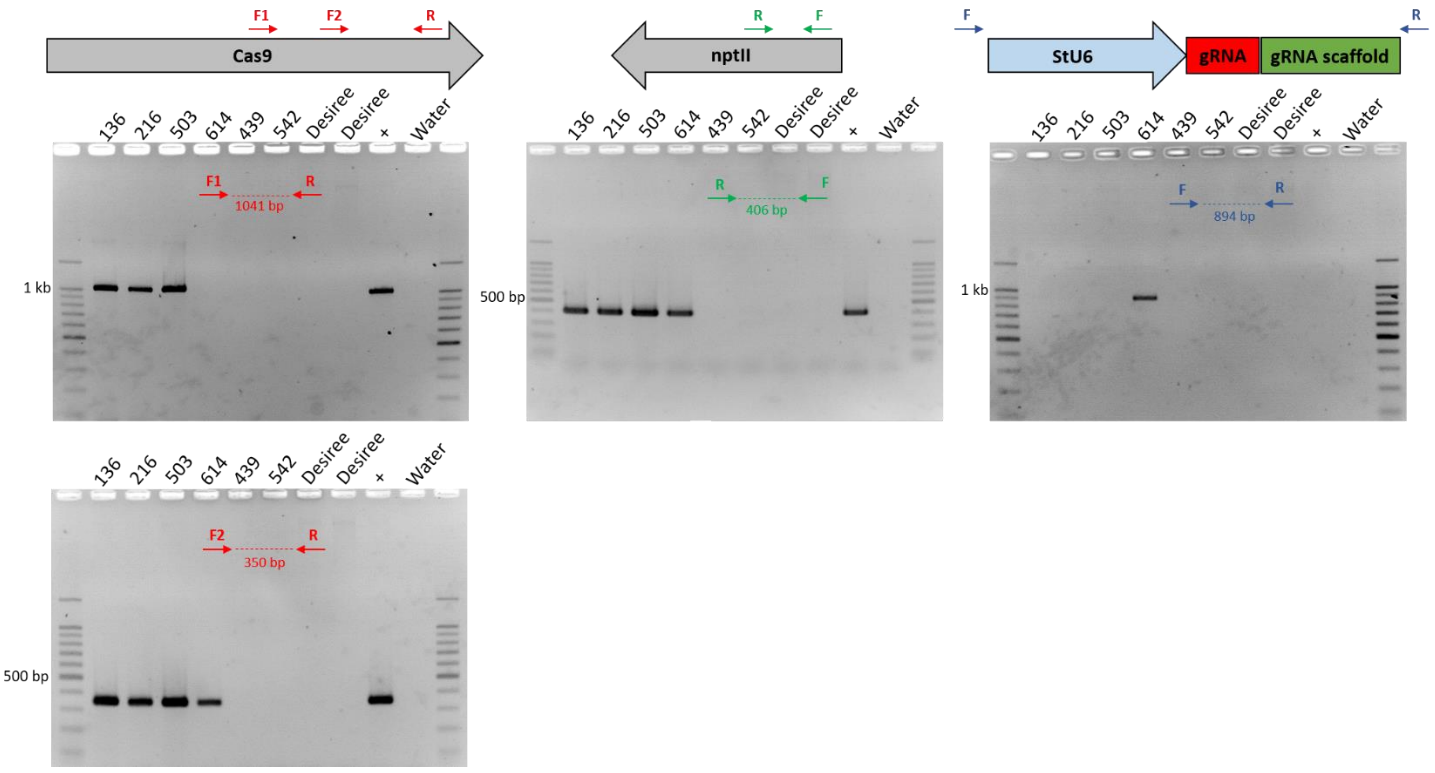

**Supplementary Fig. S7: Pictures of mutants (regenerated from PEG-transfected protoplasts) grown on medium containing kanamycin.** Positive control (control +: 17T.701.010 line stably transformed with *Agrobacterium*) and negative controls (Desiree) have been included.

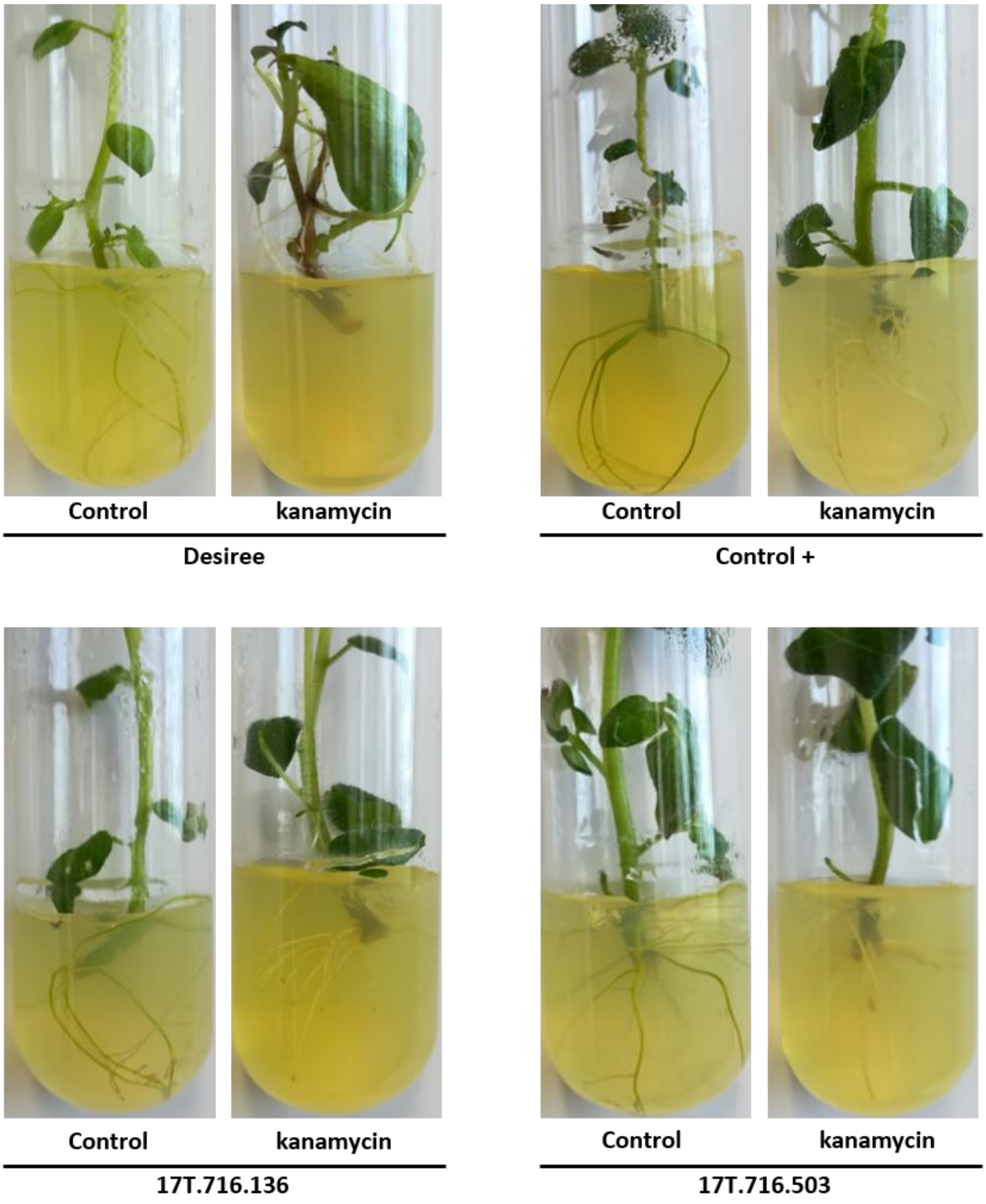

**Supplementary Fig. S8: Representative Sanger sequencing results for 5 edited lines and Desiree at the sgGBSS3-targeted site.** The PAM motif is indicated in red and the sgGBSS1-targeted locus in blue. Grey arrows indicate the location of base substitutions (C in blue, G in dark, A in green and T in red). The nature of base conversion is indicated below the chromatograms.

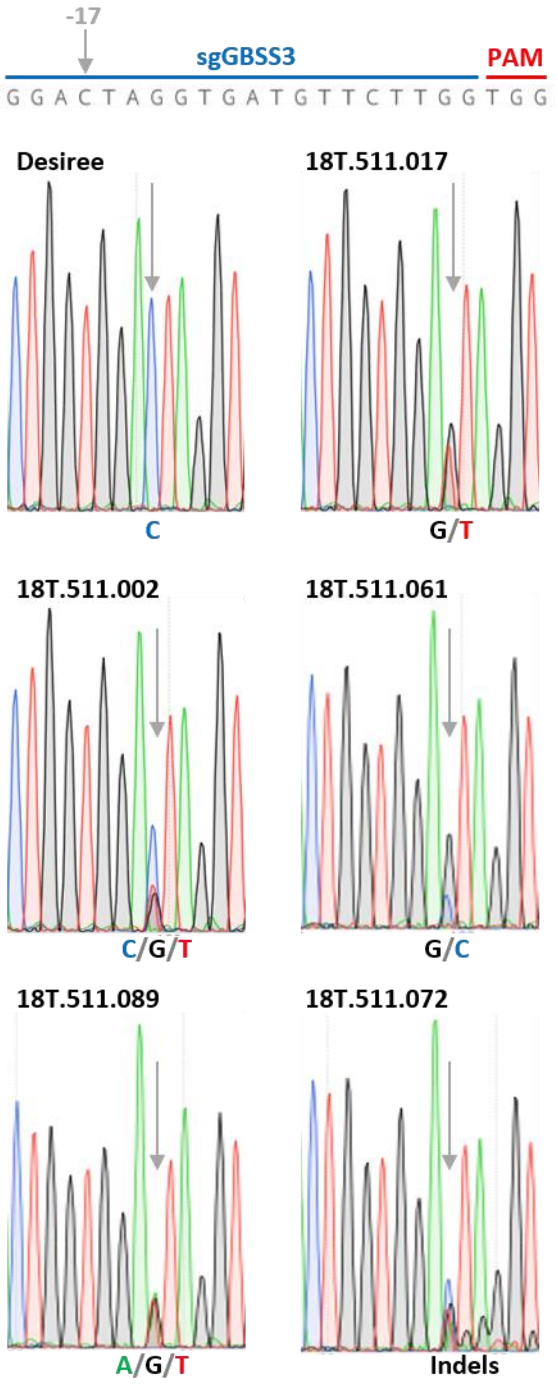
